# An Epstein-Barr virus-encoded snoRNA directs 2’-O-methylation of human rRNAs to control translation and the viral lytic switch

**DOI:** 10.64898/2026.05.13.724956

**Authors:** Yarden Dan, Beáta E. Jády, Yudit Halperin, Yuval Nevo, Aliza Fedorenko, Anat Bashan, Maram Arafat, Pnina Glick, Ada Yonath, Tamás Kiss, Daphna Nachmani

## Abstract

Epstein-Barr virus (EBV) establishes life-long latency in human B-cells yet the molecular strategies that balance its persistence with lytic replication remain incompletely understood. Here, we identify the EBV-encoded small nucleolar RNA, v-snoRNA1, as a *bona fide* 2′-O-methylation guide that directs methylation of host ribosomal RNAs at 18S-C621 and 28S-U1760, two conserved residues in the ribosomal A-site. V-snoRNA1-mediated hypermethylation impairs 18S rRNA maturation and compromises translational fidelity and output, resulting in slower cellular proliferation. Infection with a v-snoRNA1 deleted virus (Δv-snoRNA1) leads to enhanced protein synthesis and increased proliferation, together with extensive rewiring of host and viral gene expression. This rewiring includes suppression of immune and interferon pathways and alterations in transcription factor activities important for B-cell differentiation. Importantly, we find that v-snoRNA1 is required to facilitate viral production. Our findings reveal a molecular strategy by which EBV directly controls translation to promote infection.

## Introduction

The Epstein–Barr virus (EBV; also known as human herpesvirus 4 (HHV4)) is a ubiquitous human γ-herpesvirus affecting over 90% of the adult population worldwide^1^. EBV infection is etiologically linked to a range of malignancies and autoimmune disorders, including Burkitt’s and Hodgkin’s lymphomas, nasopharyngeal carcinoma, gastric carcinoma, and multiple sclerosis^1^. EBV infects both epithelial cells and B lymphocytes, where it establishes life-long latency accompanied by cellular immortalization. During latency, the viral genome persists as a nuclear episome expressing a restricted set of viral products, including the six Epstein**-**Barr nuclear antigens (EBNA1, EBNA2, EBNA3A, EBNA3B, EBNA3C, and EBNA-LP) and the three latent membrane proteins (LMP1, LMP2A, and LMP2B). These proteins promote host-cell proliferation, repress apoptosis, ensure episomal genome maintenance, and support immune evasion^2^.

Additionally, the EBV genome encodes a diverse repertoire of noncoding RNAs, including two abundant nuclear RNAs (EBER1 and EBER2), one small nucleolar RNA (v-snoRNA1), and more than 30 microRNAs^3–5^. The molecular functions of these non-coding viral RNAs are only partially understood, though they are generally believed to reinforce latency and immune evasion^6^. For example, EBER2 has been shown to base-pair with nascent transcripts from the terminal repeat region of the EBV genome and recruit the B-cell transcription factor PAX5 to modulate viral gene expression and replication^7^. This example illustrates how EBV-encoded RNAs act as molecular scaffolds that interface with host gene expression networks to sustain persistent infection. However, the functional diversity of EBV noncoding RNAs remains incompletely defined.

Ribosome biogenesis begins with nucleolar synthesis of the polycistronic 47S pre-rRNA transcript which undergoes complex nucleolytic processing, folding, and assembly of the mature 18S, 5.8S and 28S rRNAs with ribosomal proteins into small and large ribosomal subunits. These rRNAs are extensively decorated by more than 200 site-specific chemical modifications, mainly addition of 2′-O-methyl groups (2’OMe) and pseudouridine (isomerization of uridine), which are essential for accurate translation and cellular growth^8^. Perturbations in rRNA modification patterns can impair ribosome biogenesis, reduce translational fidelity, and trigger disease^9–19^. Thus, precise regulation of rRNA modification is critical for maintaining the translational capacity and function of the cell.

Site-specific 2’OMe of rRNAs is achieved by box C/D small nucleolar RNPs (snoRNPs), each comprising a sequence-specific guide snoRNA and four core proteins, including the methyltransferase fibrillarin^20–22^. Box C/D snoRNAs contain conserved C (RUGAUGA) and D (CUGA) motifs near their termini, as well as their internal imperfect copies named C′ and D′ boxes. The D or D′ box is preceded by an antisense element that base-pairs with a complementary rRNA sequence, positioning the fifth nucleotide upstream of the relevant box for methylation by fibrillarin. Besides directing rRNA 2’OMe, human box C/D snoRNAs also participate in snRNA and tRNA 2’-O-methylation^23,24^, or less frequently, in nucleolytic rRNA processing^25–28^.

EBV’s v-snoRNA1 is the only known virally-encoded snoRNA by human herpesviruses^3^, and is structurally undistinguishable from canonical box C/D 2’OMe guide snoRNAs^3,4^. However, whether it functions as a canonical snoRNA targeting rRNA, and if so, what is its physiological role during infection remained unclear. Here, we demonstrate that EBV v-snoRNA1 functions as a *bona fide* 2’OMe guide RNA with dual specificity, directing methylation of the host 18S rRNA at C621 and 28S rRNA at U1760 in a D′ and D box dependent manner, respectively. The v-snoRNA1-mediated methylation interferes with late-stage 18S processing, alters ribosomal fidelity and elongation rate, ultimately resulting in reduced global translation and in a ribosome-deficiency phenotype. During EBV infection, these methylations lead to significant rewiring of both host and viral gene expression and are required for virion production.

This work reveals a previously unrecognized strategy by which EBV directly manipulates host translation machinery through modification of rRNA and highlights viral modulation of the ribosome as a key facet in host-pathogen interactions.

## Results

### EBV v-snoRNA1 directs 2’-O-methylation of human 18S and 28S rRNAs

The EBV-encoded v-snoRNA1 exhibits all structural hallmarks of canonical box C/D 2’OMe guide snoRNAs and associates with the complete set of authentic 2′OMe snoRNP proteins^3^. Sequence complementarity analysis revealed that v-snoRNA1 is capable of base-pairing with human rRNAs at two positions (Fig. 1A). The predicted upstream antisense element (UAE) forms an 11-bp perfect helix with the G617-U627 region of 18S rRNA, positioning C621 for D′ box-dependent 2’OMe. The downstream antisense element (DAE) potentially hybridizes to the G1756–U1769 region of 28S rRNA, forming 13 bp helix interrupted by a single C–U mismatch, and is expected to direct 2′OMe at U1760.

**Figure 1.**
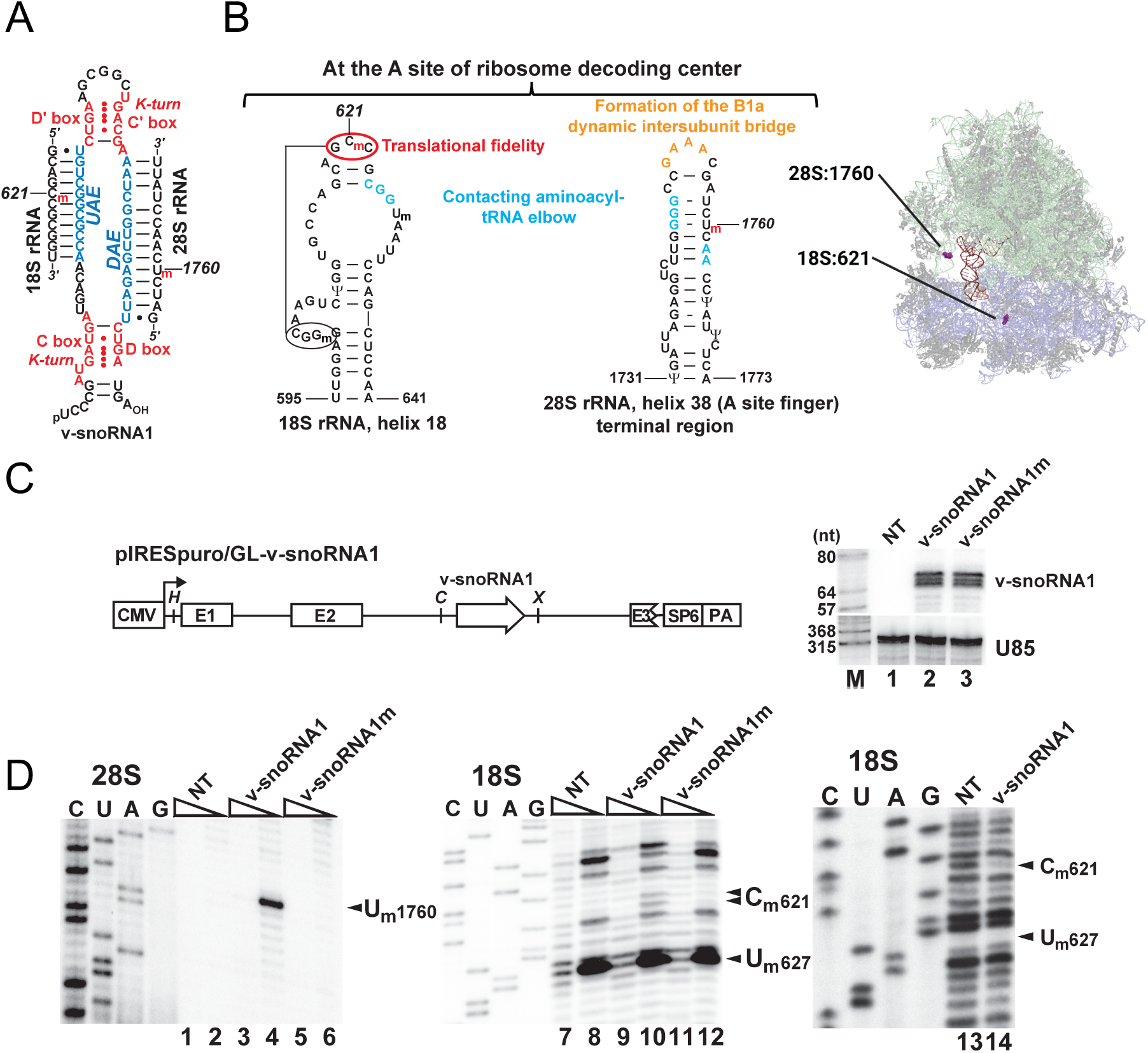
EBV v-snoRNA1 directs site-specific 2’OMe of human rRNAs. (A) Predicted two-dimensional structure of EBV v-snoRNA1 directing 2’OMe of the 18S-C621 and 28S-U1760 residues. The evolutionarily conserved C, D, C’ and D’ boxes of v-snoRNA1 forming terminal k-turn and internal k-loop structures are in red and the upstream (UAE) and downstream (DAE) antisense elements are in blue. The 2’-O-methyl groups (m) introduced by v-snoRNP1 are in red. (B) Left - Structures of the terminal hairpin regions of helix 18 of the 18S rRNA (left) and helix 38 of the 28S rRNA (right). The post-transcriptionally added 2’-O-methylated nucleotides (N_m_) are indicated. The predicted v-snoRNA1-dependent 2’-O-methyl groups at 18S-C621 and 28S-U1760 are in red. Ribonucleotides assigned for specific roles are highlighted and explained in different colors. The 18S rRNA long range interaction involving the v-snoRNA1 target nucleotide C621 is indicated. Right - The position of 18S-C621 and of 28S-U1760 on the full human ribosome structure (PDB 8glp). Light green – 28S rRNA, lilac – 18S rRNA, red – P-site, orange – mRNA, magenta – specific modified nucleotides, gray – ribosomal proteins. (C) Left - Schematic structure of the pIRESpuro/GL-v-snoRNA1 expression construct. The cytomegalovirus (CMV) promoter with the transcription start site (arrow), the exons (E1, E2, and E3), and the polyadenylation site (PA) of the human β-globin gene are indicated. The SP6 viral promoter and relevant restriction sites (*H*, *Hin*dIII, *C*, *Cla*I, and *X*, *Xho*I) are also shown. Right - RNase A and T1 mapping of ectopically expressed v-snoRNA1 and endogenous U85 scaRNA in non-transfected HeLa (NT), HeLa-v-snoRNA1 (v-snoRNA1) and HeLa-v-snoRNA1m (v-snoRNA1m) cells is shown. Lane M, DNA size markers in nucleotides. Note that mapping of v-snoRNA1 RNAs have heterogenous 5’ and 3’ termini detected by the three bands separated by 1 nt each. (D) Primer extension mapping of 2’OMe nucleotides. Terminally labeled oligonucleotide primers (complementary to the human 18S rRNA from G644 to G665 or to the 28S rRNA from G1790 to C1807 were annealed to cellular RNAs extracted from HeLa cells lacking (NT) or expressing wt or mutant EBV v-snoRNA1 and were extended with RT in the presence of 1- or 0.004-mM dNTPs as indicated above lanes 1-12. Lanes 13 and 14 show primer extension reactions performed on partially hydrolyzed template RNAs. Lanes titled C, U, A and G represent dideoxy sequencing ladders. The RT stops or gaps indicating the presence of 2’OMe nucleotides are highlighted by arrowheads on the right sides of the gels. Note that although AMV reverse transcriptase stops a nucleotide before the 2’-O-methylated nucleotides (lane 4), in some context-dependent cases, it may add a last nucleotide facing the modified nucleotide (2 stops, lane 10).

The predicted target nucleotides, 18S-C621 and 28S-U1760, reside in functionally critical regions of the ribosome^29,30^ (Fig. 1B). 18S-C621 is located in the terminal loop of helix 18 of the small subunit, adjacent to the decoding center, where helix 18 cooperates with helix 44 to monitor codon–anticodon pairing and to position ribosomal protein uS12 (RPS23) at the mRNA channel. 28S-U1760 lies within helix 38 of the large subunit, which forms the A-site finger, a structural element that contacts the elbow of the A-site tRNA, contributes to the dynamic inter-subunit bridge B1a, and supports tRNA accommodation and translocation^29,30^ (Fig. 1B). Both residues are evolutionarily conserved and lie at sites known to ensure translational fidelity and elongation, suggesting that EBV could modulate host ribosome function through v-snoRNA1-guided modification of these nucleotides.

To test whether v-snoRNA1 can direct modification of these predicted 2′OMe sites, we stably expressed wild-type v-snoRNA1 and a mutant v-snoRNA1 (v-snoRNA1m) whose UAE and DAE were replaced for complementary sequences and thus were unable to direct 2’-O-methylation, in HeLa cells using the pPIRESpuro/GL vector optimized for expression of intron-encoded snoRNAs^24,31^ (Fig. 1C). RNase A/T1 protection assays confirmed comparative accumulation of v-snoRNA1 and v-snoRNA1m in transfected cells (Fig. 1C, right). 2’OMe at 18S-C621 and 28S-U1760 was analyzed by primer extension using 5′end-labeled oligonucleotide primers (Fig. 1D). Under low dNTP conditions, AMV reverse transcriptase (RT) pauses immediately before or at 2’OMe residues^32,33^. Mapping of 28S rRNA from v-snoRNA1-expressing cells revealed a strong RT stop one nucleotide before U1760 under low dNTP conditions (Fig. 1D, lane 4), absent in control reactions performed at high dNTP or on rRNA from non-transfected or v-snoRNA1m-expressing HeLa cells (lanes 1-3 and 5-6). Similarly, mapping of 18S rRNA from v-snoRNA1 cells produced two weak but reproducible RT stops immediately before and at C621 (lane 10). A strong stop at U627 was also observed, corresponding to its known endogenous methylation by SNORD65^34^ (lanes 8, 10, 12). To verify that the observed C621 signals represented true v-snoRNA1-guided 2′OMe, primer extension was repeated on partially hydrolyzed rRNAs. Because 2′-O-methyl groups protect RNA nucleotides from alkaline hydrolysis, modified nucleotides appear as gaps in the hydrolysis ladder^20,24,32^. Indeed, 18S rRNA from v-snoRNA1-expressing cells displayed a clear protection gap at C621 (lane 14), confirming v-snoRNA1-directed methylation, albeit less efficiently than the endogenous modification at U627.

These data demonstrate that EBV v-snoRNA1 functions as a *bona fide* 2’OMe guide snoRNA, directing methylation at 18S-C621 and 28S-U1760.

### Hypermethylation of the host ribosome by v-snoRNA1 compromises 18S rRNA processing, elongation rate and global translation

By introducing methylations within conserved elements of the ribosomal A-site, we hypothesized that v-snoRNA1 might enable EBV to manipulate host ribosome activity. To determine whether the novel 2’OMe groups introduced by v-snoRNA1 affect global translation, we measured nascent protein synthesis in our HeLa cell lines. Cells expressing wt v-snoRNA1, exhibited reduced protein synthesis compared to control HeLa cells or HeLa cells expressing v-snoRNA1m (Fig. 2A), confirming that v-snoRNA1-mediated 2’OMe impairs global translation.

**Figure 2.**
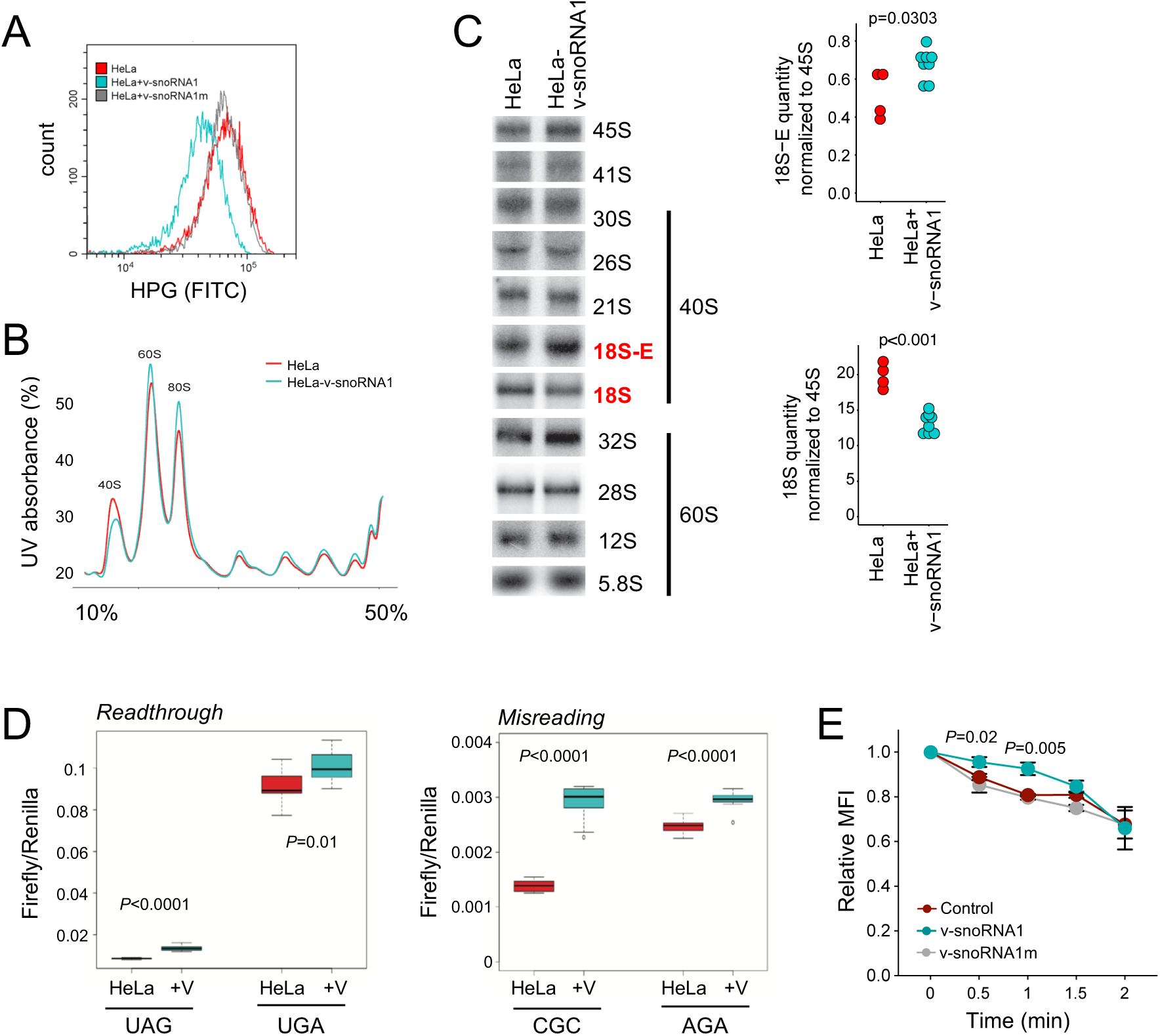
EBV v-snoRNA1 directed site-specific 2’OMe compromises cellular translation and ribosome biogenesis. (A) HeLa cells expressing v-snoRNA1 show reduced global translational activity. Nascent protein synthesis was measured in HeLa, HeLa-v-snoRNA1 and HeLa-v-snoRNA1m cells by HPG labeling. (B) Ribosome profiles obtained by sucrose gradient centrifugation. The ribosome profiles of HeLa and HeLa-v-snoRNA1 cells are drawn in red and teal, respectively. (C) Left - Steady-state accumulation of rRNA processing intermediates and mature rRNAs. Precursor and mature rRNAs were detected by Northern blot analyses. Precursor rRNA intermediates dedicated for production of the 18S small (40S) or 5.8S and 28S large (60S) ribosomal subunit rRNAs are indicated. RNAs with increased or reduced accumulation in HeLa-v-snoRNA1 cells are highlighted in red. Right – quantification of the relative abundance of 18S-E and 18S rRNAs measured in 4 independent knockout cell lines with 2 technical repeats and tested with t-test (18S-E: mean increase of 30%, p=0.0303;18S: mean decrease of 32%, p<0.001) (D) HeLa and HeLa-v-snoRNA1 overexpressing cells (+V) cells were transfected with dual luciferase reporter expressing a Renilla and firefly luciferase fusion protein. Left - For measuring readthrough, the Firefly coding sequence carried premature in frame UAG or UGA stop codons. Right - For measuring misreading, the CAC (His245) codon of the active center of firefly luciferase was replaced for CGC or AGA codon, both encoding for arginine and disrupting luciferase activity. To evaluate readthrough and misreading errors, the ratios of firefly and Renilla luciferase activities were calculated. 12 independent experiments performed, significance was calculated by t-test. (E) v-snoRNA1 expression leads to slower translation elongation. SunRiSE analysis was performed on cells expressing either v-snoRNA1, v-snoRNA1m or a Control which is an unrelated snoRNA - snoRD127. Relative MFI calculate to T=0 of Harringtonine treatment. Three independent experiments were performed; statistical significance was calculated by t-test.

Aberrant 2’OMe is known to disrupt ribosome function, stability, and biogenesis^12,14^. Thus, to investigate whether v-snoRNA1-directed methylation affects ribosome biogenesis, we analyzed ribosomal profiles by sucrose gradient sedimentation (Fig. 2B). While polysome peaks appeared normal to slightly elevated, the 40S subunit peak was consistently reduced in v-snoRNA1-expressing cells. This profile resembles the profile of ribosome deficient cells due to alteration of the 40S small subunit^35^. In this case, deficiency could arise from interference with the processing of the 18S rRNA by 18S-C621^8^. Indeed, northern blot analysis revealed a reduction in mature 18S rRNA and an increased accumulation of its 3′-extended 18S-E precursor in v-snoRNA1-expressing cells (Fig. 2C), while earlier precursors - 30S, 26S, and 21S pre-rRNAs - were unaffected. These results indicate that v-snoRNA1 expression delays the final step of 18S maturation, specifically the conversion of 18S-E to mature 18S rRNA.

Because the v-snoRNA1-directed methylations occur at the ribosomal A-site (both at the 18S and 28S), they could potentially also impact additional traits such as translational fidelity and elongation rate^36^. To assess translational fidelity, we employed dual luciferase reporter assays that measures stop-codon readthrough and amino acid misincorporation^37^. V-snoRNA1-expressing cells exhibited increased stop-codon readthrough and misreading rates compared to control cells (Fig. 2D). To assess elongation rates, we performed SunRiSE^38^ experiments of ribosome run-off and found that v-snoRNA1-expressing cells exhibit slower elongation rates compared to control cells (Fig. 2E). These results reconcile reduced global protein synthesis with unchanged polysome profiles, as slower elongation prolongs ribosome residency on mRNAs, preserving occupancy despite declining protein synthesis.

### V-snoRNA1 expression affects translation kinetics and cell proliferation

To determine whether 18S-C621 and 28S-U1760 undergo 2′-O-methylation during EBV infection, we performed RiboMethSeq analysis, across EBV-negative and EBV-positive Burkitt’s lymphoma (BL) cell lines, as well as EBV-immortalized lymphoblastoid cell lines (LCLs) (Fig. 3A-B). In EBV-negative BL cells (BJAB and Ramos), 18S-C621 was unmethylated (MethScore=0; Fig. 3A), while 28S-U1760 displayed very low basal methylation (Fig. 3B). These low methylation levels suggest that both sites are amenable to modulation by viral factors. Additionally, these data are consistent with previous studies, and support the observation that rRNA 2’OMe can occur at sub-stoichiometric levels to regulate translation^15,39–42^.

**Figure 3.**
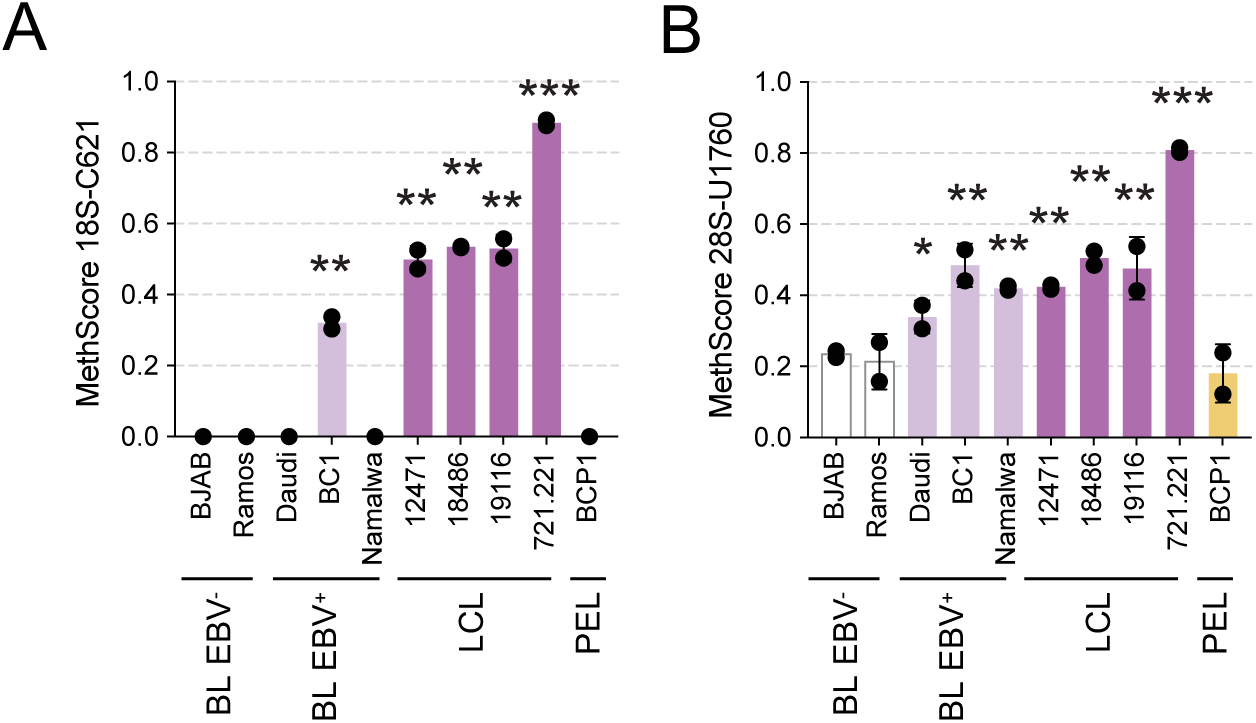
18S-C621 and 28S-U1760 are hypermethylated during EBV infection. (A, B) RiboMethSeq analysis of Burkitt’s lymphoma (BL) cell lines (either EBV-negative, or EBV-positive), of lymphoblastoid cell lines (LCL) immortalized with EBV, and of BCP1, a primary effusion lymphoma (PEL) cell line which is KSHV-positive and EBV-negative. (A) MethScore of 18S-C621, and (B) MethScore of 28S-U1760 in the above-mentioned cells. MethScore of 0 represents no methylation and 1 represents complete methylation. Two repeats were performed, and statistical significance was calculated by t-test. *-*P*<0.01, **-*P*<0.001, compared to methylation in BJAB.

In EBV-positive BL cell lines, we found that 28S-U1760 was hypermethylated, whereas 18S-C621 methylation was more restricted and detected only in BC1 cells (Fig. 3A-B, light purple bars). In LCLs both sites were hypermethylated (Fig. 3A-B, dark purple bars). This methylation pattern appeared to be specific to EBV-infected cells, as primary effusion lymphoma cells infected with Kaposi’s sarcoma-associated herpesvirus, but not with EBV, displayed methylation levels comparable to those of EBV-negative BL controls (Fig. 3A-B, BCP1, orange bar).

To assess the contribution of v-snoRNA1 to 2′-O-methylation in B cells independently of viral infection, we overexpressed v-snoRNA1 in the EBV-negative BL cell line BJAB, providing a physiologically relevant cellular context. Expression of wild-type v-snoRNA1, v-snoRNA1m, and an unrelated control snoRNA was confirmed (Fig. S1A), along with the corresponding 2′-O-methylation levels (Fig. 4A, S1B).

**Figure 4.**
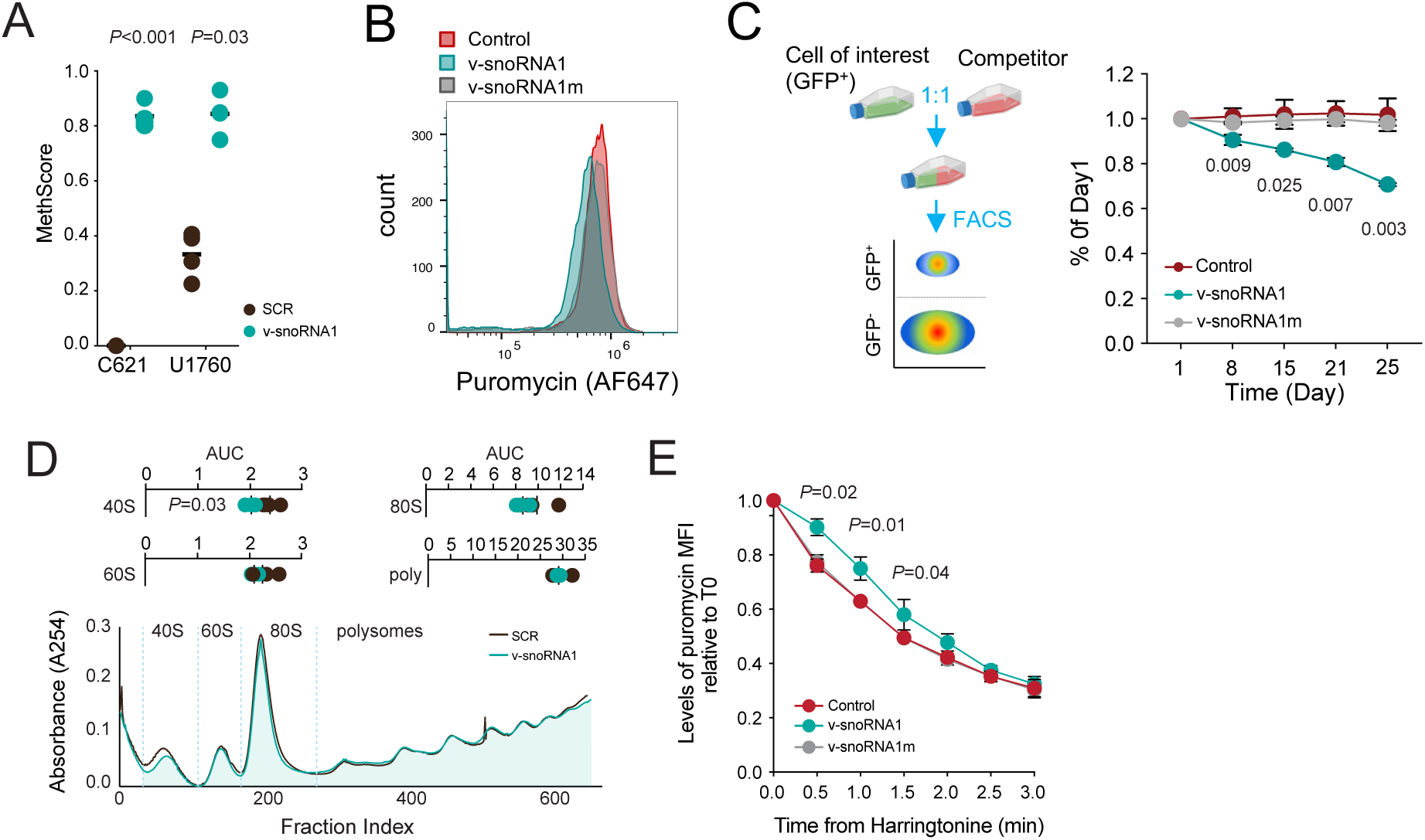
Ectopic expression of v-snoRNA1 leads to reduced global translation, elongation rate and cell proliferation. (A) MethScore of 18S-C621 and 28S-U1760 in BJAB cells stably expressing either the scrambled sequence of v-snoRNA1 (SCR) or v-snoRNA1. (B) BJAB cells expressing v-snoRNA1 (teal) show reduced global translational activity by puromycin incorporation tested in flow cytometry. Control – an unrelated snoRNA, snoRD127. (C) v-snoRNA1-expressing cells show reduced proliferation. Left - illustration of competitive proliferation assay. Cells overexpressing v-snoRNA1, v-snoRNA1m or control are GFP^+^, and were mixed at a ratio of 2:1 with GFP^-^ BJAB competitors. The proportion of each cell in the population was quantified by flow cytometry. Right – results of competitive assay. The y-axis is the percentage of each population relative to its proportion at Day 1 (day of initiation of competition). Three biological repeats were performed; statistical significance was calculated by t-test. (D) Ribosome profiles obtained by sucrose gradient centrifugation. The ribosome profiles of cells expressing a scramble sequence of v-snoRNA1 (SCR) and v-snoRNA1-expressing BJAB cells are drawn in black and teal, respectively. Dashed lines mark the borders of each fraction. To calculate significance, the area under the curve (AUC) was calculated for 3 independent experiments. (E) v-snoRNA1 expression leads to slower translation elongation. SunRiSE experiments were performed on cells expressing either v-snoRNA1, v-snoRNA1m or an unrelated snoRNA - snoRD127. Relative MFI calculate to T=0 of Harringtonine treatment. Three independent experiments were performed; statistical significance was calculated by t-test.

Analysis of global translation revealed that, as observed in HeLa cells (Fig. 2A), v-snoRNA1 overexpression in BJAB cells led to a decrease in global translation (Fig. 4B). Given the well-established link between translation and cell proliferation^43,44^, we assessed the proliferative capacity of v-snoRNA1-overexpressing cells using a competitive proliferation assay (Fig. 4C, left). V-snoRNA1 overexpression resulted in reduced cellular proliferation (Fig. 4C, right). This effect was dependent on both methylation-sites as expression of mutant v-snoRNA1 variants, either single-site or double mutants incapable of directing 2’OMe (Fig S1C), failed to suppress proliferation (Fig. 4C right, Fig. S1D).

At the molecular level, similar to our observations in HeLa cells (Fig. 2B), the v-snoRNA1-mediated decreased global translation (Fig. 4B) was accompanied by a decrease in the 40S fraction only (Fig. 4D). This pattern has been reported as a feature of ribosome-deficiency states^35^. SunRiSE experiments demonstrated that also in BJAB cells, elongation was significantly slower in v-snoRNA1-overexpressing cells compared to cells expressing v-snoRNA1m or an unrelated snoRNA control (Fig. 4E).

### V-snoRNA1 regulates global translation and proliferation during EBV infection

To validate the function of v-snoRNA1-mediated rRNA methylation during EBV infection, primary B-cells were infected with either wild-type (WT) EBV of the M81 strain^45,46^ or a M81 mutant lacking v-snoRNA-1^3^ (Δv-snoRNA1) thus generating new LCLs (Fig. S2A). RiboMethSeq analysis showed that deletion of v-snoRNA1 restored methylation at both 18S-C621 and 28S-U1760 to the basal levels observed in EBV-negative BL cells (Fig. 5A and Fig. 3, respectively).

**Figure 5.**
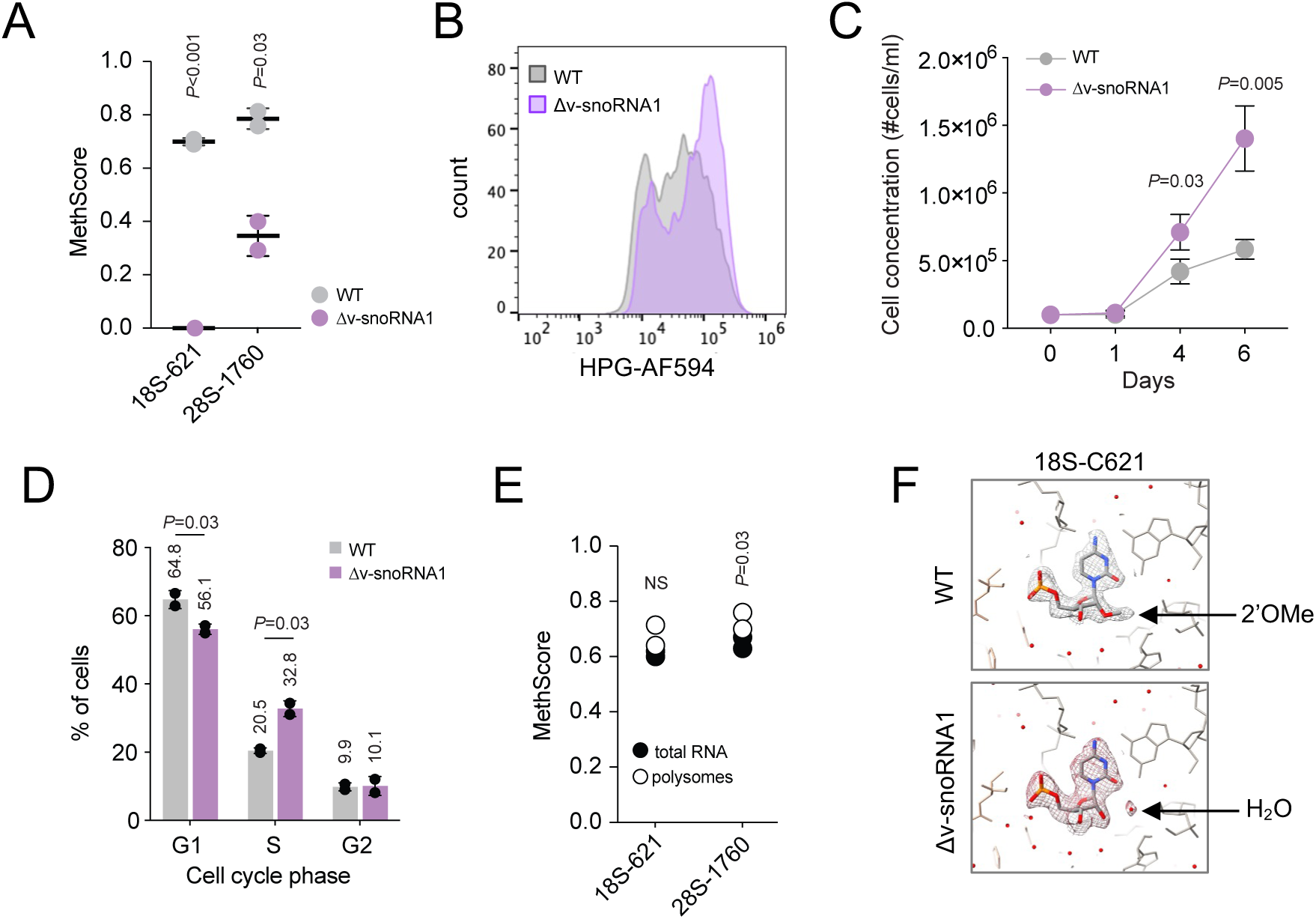
During EBV infection, v-snoRNA1 is functional in regulating translation and proliferation. (A) RiboMethSeq analysis was performed on LCLs infected either with WT or Δv-snoRNA1 of the M81 EBV strain. Presented are the MethScores of 18S-C621 and of 28S-U1760. Statistical significance was calculated by t-test. (B) Nascent protein synthesis analysis of EBV WT- and Δv-snoRNA1-infected LCLs. Δv-snoRNA1 exhibited higher incorporation of the methionine analogue L-Homopropargylglycine (HPG). (C) Cell proliferation of EBV WT- and Δv-snoRNA1-infected cells was assessed by cell counting at the indicated time points, initiated at a concentration of 100,000 cell/ml. Five biological repeats were performed and statistical significance was calculated by t-test. (D) Cell cycle analysis of WT- and Δv-snoRNA1-infected cells, depicting a skewing in the cell cycle of Δv-snoRNA1-infected cells relative to WT-infected cells. Three biological repeats were performed, and statistical significance was calculated by t-test. (E) RiboMethSeq of polysome fraction of EBV WT-infected LCLs, demonstrating that hypermethylated ribosomes are actively translating ribosomes. (F) EM density surrounding 18S-C621 in ribosome structures of polysomes purified from LCLs infected either with WT (top) or with Δv-snoRNA1 of the M81 EBV strain (bottom). Ribosomes from WT-infected cells show clear 2’OMe on 18S-C621, while Δv-snoRNA1 infected cells show a water molecule occupying the freed space.

We next investigated the molecular and cellular consequences of v-snoRNA1-mediated 2’-O-methylation during EBV infection. Δv-snoRNA1-infected LCLs showed the opposite phenotype to v-snoRNA1 ectopic expression – these cells exhibited increased global protein synthesis (Fig. 5B), and had a higher proliferation rate compared with WT-infected cells (Fig. 5C; Fig. S2B). Consistent with this, cell-cycle analysis revealed a reduced proportion of cells in G1 phase and an increased proportion in S phase in Δv-snoRNA1-infected cells relative to WT-infected cells (Fig. 5D).

To better understand the structural consequences of these modifications, we performed structural analyses of polysomes isolated from WT– and Δv-snoRNA1–infected LCLs. We first carried out RiboMethSeq analysis of the polysome fractions, confirming that hypermethylated ribosomes are actively engaged in translation (Fig. 5E). Our structural analyses revealed clear density consistent with a methyl group at 18S-C621 in WT-infected cells, which was replaced by an ordered water molecule (or possibly a potassium ion) in Δv-snoRNA1-infected cells (Fig. 5F). This water molecule forms a network of contacts with neighboring 18S residues, specifically with the O2′ and O2 of C621, the O4′ of G7, the N3 and O2′ of G6, and the O4′ of C622 (Fig. S2C), likely contributing to local stabilization of this region of helix 18, although it is unlikely to be the sole stabilizing element. On the opposite face of this water-mediated network lies ribosomal protein uS12 (RPS23), which, in turn, makes direct contacts with the mRNA at the decoding center. Notably, the phosphate oxygens of C621 form direct interactions with the backbone amide of Gly113 and the side chain of Arg107 of uS12, while the contact between the mRNA and uS12 occurs with the side chain of Gln61 and Pro62 (Fig. S2D). Exchange between 2’OMe and a water molecule at C621, and the consequent possible difference in stability around this position, may therefore propagate to uS12 and subtly alter its positioning at the mRNA-decoding interface. Methylation at 28S-U1760 places a 2′OMe group at a functionally pivotal interface, where it could potentially influence A-site tRNA docking, accommodation kinetics, or A-site finger flexibility^29^. Unfortunately, 28S-U1760 could not be resolved at sufficient resolution to model directly, due to local conformational flexibility in this region of the A-site finger ^29^.

Taken together, these findings demonstrate that during infection, v-snoRNA1 guides 2’OMe of host rRNAs at the ribosomal A-site. These site-specific modifications result in ribosome-deficient phenotype which includes decreased translation elongation kinetics, reduced protein output, and a decline in proliferation rates.

### Rewiring of host gene-expression networks by v-snoRNA1

Ribosome deficiencies are known to have widespread impact on gene expression^35,47–49^. Thus, to further explore the molecular consequences of v-snoRNA1 expression and ribosomal hypermethylation, we performed ribosome profiling (which include both RNA-seq and sequencing of ribosome protected fragments (RPF) of the same samples), and proteomic profiling of EBV WT- and Δv-snoRNA1-infected LCLs. These analyses revealed widespread and significant alterations to host transcriptome, translatome and protein abundance (Fig. 6A). Principal component analysis (PCA) demonstrated clear separation between WT and Δv-snoRNA1 samples, across all levels of analysis, underscoring extensive remodeling of the host gene expression (Fig. S3A). These datasets also showed relatively high inter-dataset correlations (Fig. S3B). Gene set enrichment analysis (GSEA) revealed strong positive and negative common enrichment across datasets (Fig. 6B; Fig. S3C). Specifically, immune- and interferon-related pathways, as well as metabolic processes, were negatively enriched in Δv-snoRNA1-infected cells (Fig. 6B). In contrast, pathways associated with ribosome biogenesis, the cell cycle and RNA processing were positively enriched (Fig. 6B). Interestingly, despite the significant enrichments to ribosome-related pathways, ribosomal protein abundance remained unaltered (Fig. S3D), further indicating that differences in translational capacities between Δv-snoRNA1-infected and WT-infected cells likely arises from altered ribosomal function rather than changes in ribosome quantity.

**Figure 6.**
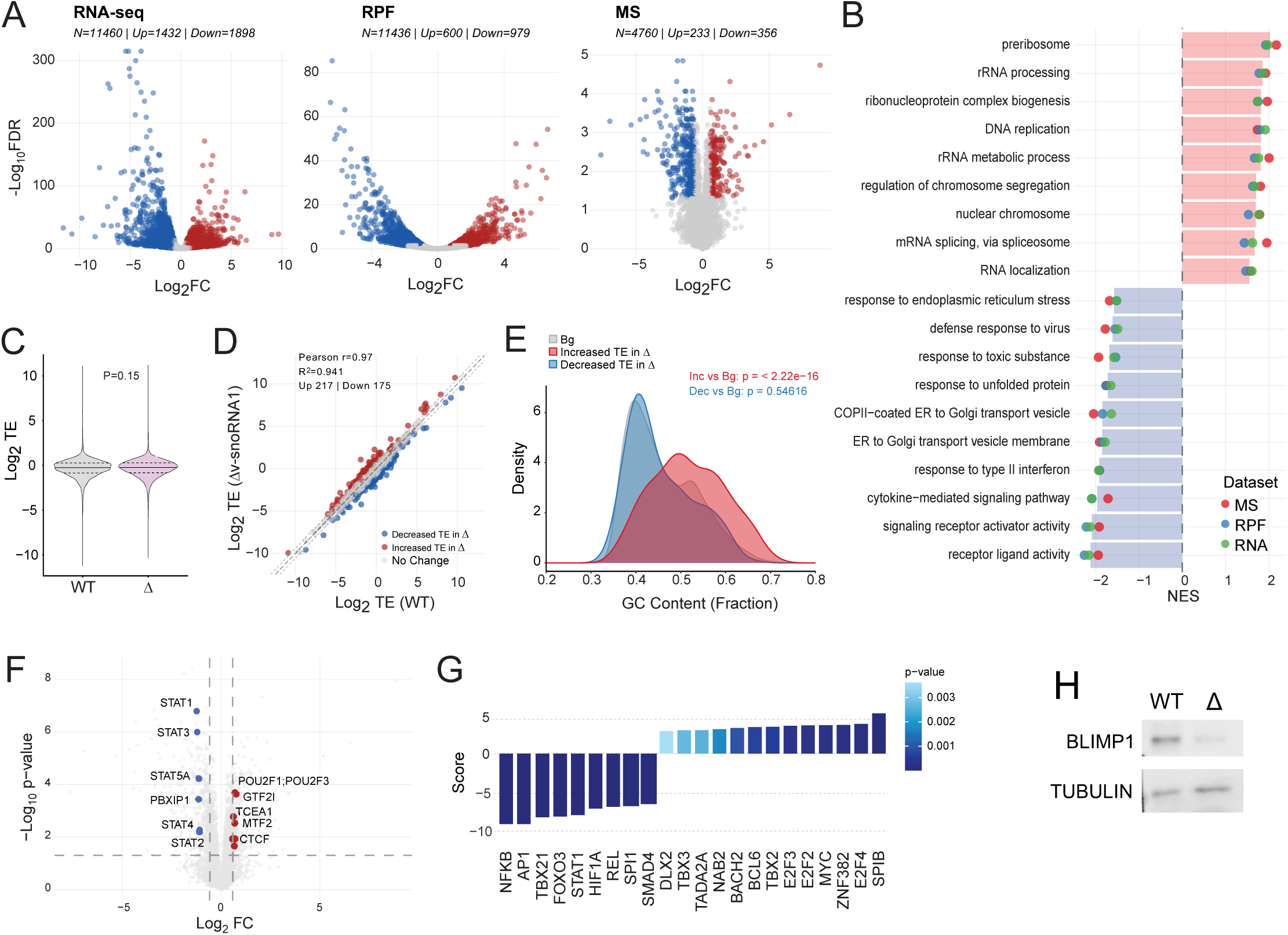
V-snoRNA1 expression leads to a widespread change in host gene expression programs. (A) Volcano plots of Log2 fold change of Δv-snoRNA1-infected cells versus WT-infected cells. Presented are datasets of RNA-seq, ribosome protected fragments sequencing (RPF) and proteomic profiling. In blue, significantly reduced proteins in Δv-snoRNA1-infected cells. In red, significantly increased proteins in Δv-snoRNA1-infected cells. (B) Top common pathways identified by GSEA in RNA-seq, RPF and proteomics, comparing Δv-snoRNA1-infected cells versus WT-infected cells. (C) Global TE analysis of WT- and Δv-snoRNA1-infected cells showed no significant differences in global TE distributions. Statistical significance was calculated with Wilcoxon test. (D) Correlation of TE values of WT-infected and Δv-snoRNA1-infected cells. Dotted lines represent the 0.58 threshold of differential TE. (E) Transcripts with increased TE (threshold of 0.58) in Δv-snoRNA1-infected cells have a higher GC-content across their full transcript sequence. Statistical significance was calculated with Wilcoxon test. (F) Volcano plot of proteomics analysis showing specific transcription factor abundance in Δv-snoRNA1-infected cells versus WT-infected cells. In blue, significantly reduced proteins in Δv-snoRNA1-infected cells. In red, significantly increased proteins in Δv-snoRNA1-infected cells. (G) Transcriptional activity analysis based on the differential expression of transcription factor target genes. Transcriptional programs that significantly differed between WT- and Δv-snoRNA1-infected cells are presented. Negative values represent reduced activity in Δv-snoRNA1-infected cells while positive values represent increased activity in Δv-snoRNA1-infected cells. (H) western blot of BLIMP1 in EBV WT- and Δv-snoRNA1-infected cells (WT, and Δ, respectively). TUBULIN served as a loading control.

To gain additional insight into ribosomal translational capacity in Δv-snoRNA1-infected versus WT-infected cells, we performed a translation efficiency (TE) analysis of our ribosome profiling dataset. Global TE remained comparable between Δv-snoRNA1-infected and WT-infected cells (Fig. 6C), with 217 genes showing increased TE and 175 with decreased TE (Fig. 6D). Sequence analyses revealed that genes with increased TE in Δv-snoRNA1-infected cells possess a higher GC content throughout the full length of their transcripts (Fig. 6E). GC content serves as an important determinant of RNA secondary structure, which plays a central role in determining TE under normal conditions and in ribosome-deficiencies^35,50^. Thus, the finding of increased TE of transcripts with high GC contents in Δv-snoRNA1-infected cells aligns well with the finding that the EBV-mediated methylations affect 40S abundance, but not of polysomes (Fig. 4D), and alters translation elongation rate (Fig. 4E).

Given the extensive gene expression remodeling, we hypothesized that changes originating from altered translation might, as a secondary effect, impact transcription. Indeed, we found changes in the protein abundance of several transcription factors, including reduced levels of several STAT proteins important for B-cell biology^51^ (Fig. 6F). To further assess the impact on transcriptional programs, we performed transcriptional network analysis of our RNA-seq dataset, further revealing perturbations in key regulatory programs, including those associated with transcription factors whose protein levels changed (*e.g. STAT1*), as well as additional major regulators of B-cell differentiation and proliferation - *SPI1*, *MYC*, *BACH2*, *BCL6*, and *SBIP*^52^ (Fig. 6G). For example, *BACH2*, which promotes B-cell differentiation to plasma cells by repressing the transcription factors *BLIMP1* and *XBP1*^53–57^, exhibited increased activity in Δv-snoRNA1-infected cells compared to WT. Consistently, transcript levels of *PRDM1* (encoding the BLIMP1 protein) and *XBP1* were markedly reduced (Log₂FC = –2.9, adjusted p = 1.21×10⁻¹²⁹; and Log₂FC = –1.08, FDR = 5.15×10⁻²², respectively). While not detected in our proteomics profiling, western blot analysis confirmed substantial decrease in BLIMP1 protein abundance in Δv-snoRNA1-infected cells (Fig. 6H), consistent with enhanced *BACH2* activity.

Collectively, these results demonstrate that v-snoRNA1 expression leads to a widespread impact on gene expression which governs B-cell identity during EBV infection.

### Viral production requires hypermethylated ribosomes

Due to the significant gene expression changes of the host, we now turned to analyze the impact of v-snoRNA1 on viral gene expression. Our proteomic profiling identified only a limited number of viral proteins (Fig. 7A). Among these, the lytic proteins BMRF1, MCP, BFRF1, and BALF2 were markedly reduced in Δv-snoRNA1-infected cells, whereas EBNA1, a key latency-associated factor, was upregulated. Our RNA-seq and RPF data showed clear differences in viral gene expression between WT- and Δv-snoRNA1-infected cells (Fig. 7B-C). RNA-seq data showed that in Δv-snoRNA1-infected cells, latency-associated transcripts such as *EBNA1, EBNA2,* and *EBNA3B/C* were highly abundant, while lytic regulators, including *BZLF1*^58,59^ and *BMRF1*^60^, were reduced (Fig. 7B). RPF data show decreased ribosome occupancy on lytic transcripts and enhanced occupancy on latent genes, including *EBNA3B/C* (Fig. 7C), in Δv-snoRNA1-infected cells. Calculation of global TE of viral transcripts revealed reduced TE distributions in Δv-snoRNA1-infected cells (Fig. 7D). Notably, this effect was specific to lytic transcripts, as latent transcripts displayed comparable TE distribution between WT- and Δv-snoRNA1-infected cells (Fig. 7E-F). Unfortunately, the limited number of viral genes did not enable us to perform sequence-based analysis to identify shared motifs or regulatory elements underlying this selectivity.

**Figure 7.**
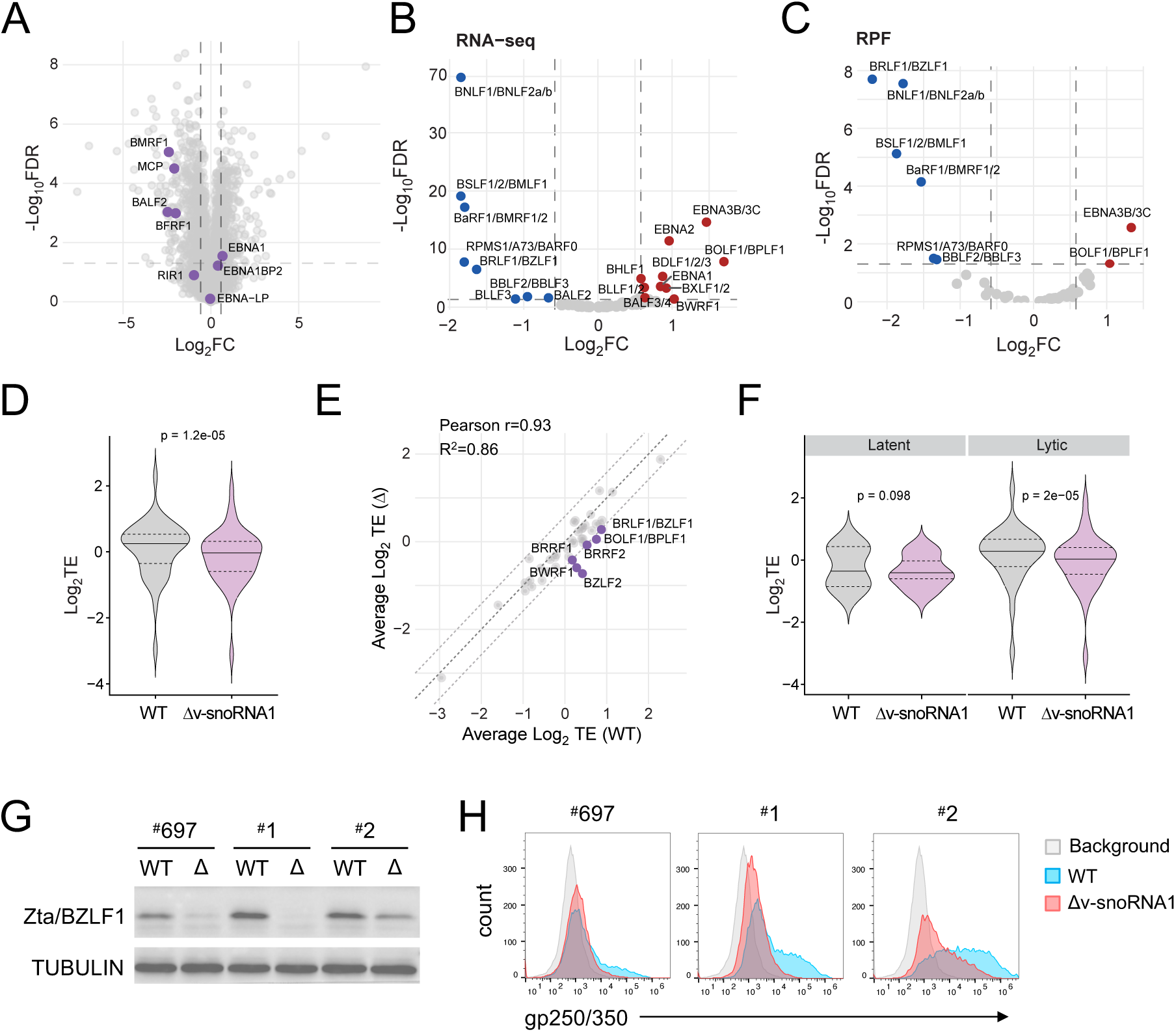
v-snoRNA1 regulates the viral proteome and lytic switch. (A) Volcano plot of proteomic profiling highlighting viral proteins which were detected in WT-and Δv-snoRNA1-infected samples. Viral proteins are marked in purple; host proteins are marked in grey. Positive values represent increased abundance in Δv-snoRNA1-infected cells, negative values represent decreased abundance in Δv-snoRNA1-infected cells. (B) Differential expression of viral transcripts by RNA-seq analysis. In blue, significantly reduced transcripts in Δv-snoRNA1-infected cells. In red, significantly increased transcripts in Δv-snoRNA1-infected cells. (C) Differential ribosome occupancy of viral transcripts (ribosome protected fragments). In blue, significantly reduced transcripts in Δv-snoRNA1-infected cells. In red, significantly increased transcripts in v-snoRNA1-infected cells. (D) TE values of viral transcripts only in WT- and Δv-snoRNA1-infected samples. Significance calculated by Paired Wilcoxon test. (E) Correlation of TE. Dotted lines represent the 0.58 threshold of differential TE. (F) TE of either latent or lytic transcripts in WT- or Δv-snoRNA1-infected samples. Significance calculated by Paired Wilcoxon. (G) western blot analysis of viral BZLF1 in three additional LCLs infected with WT and Δv-snoRNA1 viruses. Number at the top indicate internal numbering of specific donor. (H) FACS detection of the virion gp250/350 glycoprotein of WT- and Δv-snoRNA1-infected cells (blue and red histograms respectively). The grey histogram is the background staining by secondary antibody only.

In light of the above findings of differential TE of latent and lytic genes, we next focused on *BZLF1*, a master regulator of the latent-to-lytic switch^58,59^, which showed reduced abundance in Δv-snoRNA1-infected cells based on both RNA-seq and RPF frequencies and TE values (Fig. 7B-C and E). This reduction suggests that v-snoRNA1 may be required to sustain basal BZLF1 expression, thereby facilitating productive viral replication. Western blot analysis of LCLs from additional donors confirmed robust BZLF1 downregulation in Δv-snoRNA1-infected LCLs (Fig. 7G). To assess the functional outcome of the differential gene expression of Δv-snoRNA1-infected cells, we quantified viral production by flow cytometric detection of the envelope glycoprotein gp250/350. Δv-snoRNA1-infected cells displayed minimal gp250/350 staining, pointing to impaired virion production (Fig. 7H), which is consistent with low BZLF1 levels (Fig. 7G).

Taken together, these results suggest a role for v-snoRNA1 in sustaining the latent-lytic balance necessary for viral propagation.

## Discussion

EBV has evolved diverse strategies to manipulate the host cell to its own benefit^6,61–64^. Here, we identify the EBV-encoded v-snoRNA1 as a true guide for 2’OMe of two residues in host rRNAs. Our findings reveal a previously unrecognized layer of viral manipulation, whereby EBV tailors the ribosome itself to control cellular and viral gene expression during infection with implications on viral production.

V-snoRNA1 was first identified by Hutzinger *et al.*^3^, who showed that it associates with canonical snoRNP proteins in the EBV B95.8 strain; however, no methylation target was identified. Here, using primer extension, RiboMethSeq and structural analyses, we demonstrate that v-snoRNA1 guides 2’OMe at 18S-C621 and 28S-U1760, two residues located within the ribosomal A-site. We find that hypermethylated ribosomes are deficient in translational capacity, exhibiting slow elongation, reduced fidelity and a defect in ribosome biogenesis. Specifically, methylation at 18S-C621 impairs the final maturation step of 18S rRNA - the NOB1-mediated cleavage of 18S-E to mature 18S (Fig. 2C). As NOB1 activity depends on the assembly factor Rio1, which engages helix 18 within pre-40S particles^65,66^, 2′OMe of C621 which is located in the terminal loop of helix 18, might interfere with Rio1 recruitment or NOB1-mediated cleavage, thereby delaying 18S-E processing. Our structural analyses also suggest a role for methylation of 18S-C621 in regulating translation fidelity. As the methyl group at 18S-C621 occupies a position otherwise filled by a water molecule, the methyl group might subtly reposition uS12 (Fig. 5F, S2C-D) and influence decoding fidelity, consistent with the reduced fidelity observed in hypermethylated ribosomes (Fig. 2D). Additionally, the reduced elongation rate might be attributed to methylation of 28S-U1760, as this residue resides in the A-site of the large ribosomal subunit, a region in close proximity to the D-loop of the A-site tRNA and to elongation factor binding interfaces^67–69^.

To investigate the cellular impact of v-snoRNA1 expression during EBV infection, we reasoned that, given the long half-life of ribosomes^70,71^, a prolonged infection period would be required to allow the turnover of pre-existing unmodified ribosomes and the accumulation of newly assembled, hypermodified ribosomes. We therefore examined later stages of infection using newly established LCLs from B cells infected with either WT or Δv-snoRNA1 viruses of the M81 strain, which displays higher lytic activity than the previously studied B95.8 strain^3,45,46^. In these settings, we found that v-snoRNA1 contributes to sustaining viral production capacity.

Our findings suggest two complementary paths by which v-snoRNA1 might rewire the host cell and control viral production. In the first, host-centered pathway, v-snoRNA1-mediated molecular rewiring originates at the ribosome, leading to differential host protein expression that progressively propagates through cellular networks. Notably, the functional state induced by v-snoRNA1-mediated hypermethylation closely resembles ribosome deficiency, a condition in which reduced ribosomal capacity not only reshapes the translatome but also exerts profound secondary effects on transcription, as demonstrated in several model systems^35,47,48,72,73^. Consistent with this model, as infection progresses, the primary translational perturbations are amplified through downstream effects on transcription factors and other regulatory networks, ultimately reshaping cellular gene expression landscapes with consequences for viral infection. Indeed, deletion of v-snoRNA1 altered key transcriptional circuits during infection - for example, by leading to elevated BACH2 transcriptional program, which in turn suppressed the expression of BLIMP1 and XBP1, factors required for B-cell differentiation to plasma cells^53–56^. Since the EBV lytic switch, BZLF1, is induced during B-cell differentiation^74^, this impaired transcriptional program may underlie the reduced BZLF1 expression and diminished viral production observed in Δv-snoRNA1-infected cells.

In the second, more virus-centered pathway, v-snoRNA1 is required for the translation of viral transcripts (Fig. 7B-F). Given our molecular findings that v-snoRNA1-mediated methylations decrease translational fidelity (Fig. 2D), and lead to a translational bias of viral transcripts (Fig. 7D-F), it is plausible that hypermethylated ribosomes are required to facilitate basal translation of EBV lytic transcripts, thereby fine-tuning the balance between latency and lytic replication. Interestingly, in this context, recent studies from the Whitehouse laboratory demonstrated that KSHV, a close relative of EBV, manipulates ribosome assembly and post-transcriptional modification to promote translation of lytic genes from upstream open reading frames, and showed this mechanism is essential for virion production^75,76^. Whereas KSHV achieves ribosome manipulation by targeting host proteins, our findings reveal that EBV employs an alternative strategy - encoding a viral snoRNA that directly guides modification of the host ribosome. Together, these observations suggest that both KSHV and EBV have evolved distinct yet convergent tactics to manipulate host ribosomes towards the same end - control of virion production. This convergence mirrors the microRNA-mediated immune evasion strategies of these viruses, in which EBV and KSHV each encode a unique miRNA targeting a common immune ligand to evade immune recognition^77^.

Lastly, our findings position v-snoRNA1 as a potential therapeutic agent for modulating both viral infection and host cell proliferation. Expression of this viral snoRNA, irrespective of viral context, impairs translation and proliferation of uninfected cells (Fig. 4C), suggesting that it may also hold therapeutic potential for proliferative disorders susceptible to ribosome dysfunction, independent of EBV infection.

In summary, our results highlight the diversity of viral mechanisms for host manipulation. They underscore the emerging importance of rRNA modifications and ribosome modulation for physiological processes including viral infection.

## Resource Availability

### Lead Contact

Requests for further information and resources should be directed to and will be fulfilled by the lead contact, Daphna Nachmani.

### Materials Availability

Plasmids and cell lines generated in this study will be made available on request, but we may require a payment and/or a completed materials transfer agreement if there is potential for commercial application.

### Data and Code Availability

Ribo-seq and RNA-seq data supporting the findings of this study are available in the NCBI Sequence Read Archive (SRA) repository, under the BioProject accession number PRJNA1359357. Proteomics data is available as Supplementary Material. Ribosome structures were deposited at PDB - WT: 9TT7, EMD-56227, delta: 9TV2, EMD-56286.

## Acknowledgments

DN is supported by the European Research Council (10107560). DN Received support from the Abisch-Frenkel Faculty Development Lectureship and by the Alon Fellowship from the Israeli Council for Higher Education. DN is a member of the Human RNome Consortium. TK was supported by a grant from l’Agence National de la Recherche (ANR-22-CE12-0020). We thank Henri-Jacques Delecluse for kindly providing all EBV M81, WT- and Δv-snoRNA1-infected LCLs. We are grateful to Professor Erik C. Boettger and Dr. Harshita Santosh Kumar for providing us the pRM hRluc-hFluc vectors. We thank Dr. Marie-Françoise O’Donohue for providing us oligonucleotides and advice for the detection of ribosomal precursors. We acknowledge the use of BioRender.com for creating illustrations.

## Author Contributions

B.E.J and D.N designed research. Y.D, B.E.J and D.N performed research. Y.H. and Y.N. performed computational analyses. A.F., A.B. and A.Y solved ribosomal structure. M.A and P.G, provided vital assistance. B.E.J, T.K and D.N. wrote the manuscript. T.K and D.N supervised the project.

## Declaration of Interests

All authors declare no conflict of interest

---

**Supplementary Figure 1.**
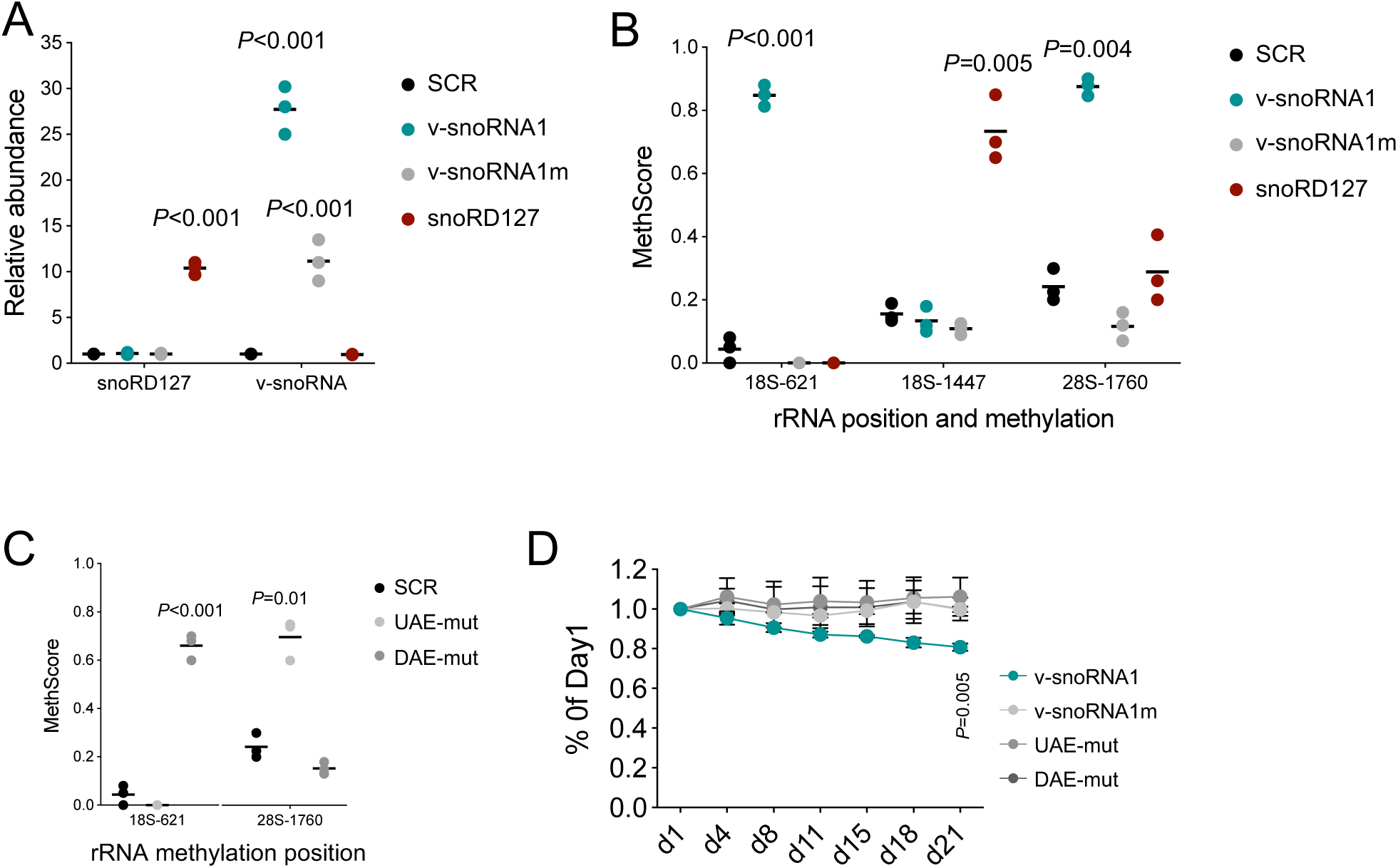
Expression of v-snoRNA1 in BJAB cells. Related to. Figure 4. (A) qPCR validation of expression of v-snoRNA1, v-snoRNA1m and snoRD127. Three independent experiments were performed. Statistical significance was calculated by t-test relative to SCR (the scrambled sequence of v-snoRNA1). (B) RiboMethSeq analysis in the various cells in (A) to validate 2’OMe, showing that v-snoRNA1 facilitates methylation of 18S-C621 and of 28S-U1760 while v-snoRNA1m does not. SnoRD127 overexpression was successful in increasing methylation of 18S-1447 specifically. Three independent experiments were performed. Statistical significance was calculated by t-test, relative to SCR (the scrambled sequence of v-snoRNA1). (C) RiboMethSeq analysis demonstrating the inability of the UAE and DAE mutants to facilitate 2’OMe. (D) v-snoRNA1 leads to reduced proliferation in a 2’-*O*-methylation-dependent manner. Competitive proliferation assay of BJAB cells expressing either v-snoRNA1, v-snoRNA1m, UAE-mut or DAE-mut. Statistical significance was calculated by t-test.

**Supplementary Figure 2.**
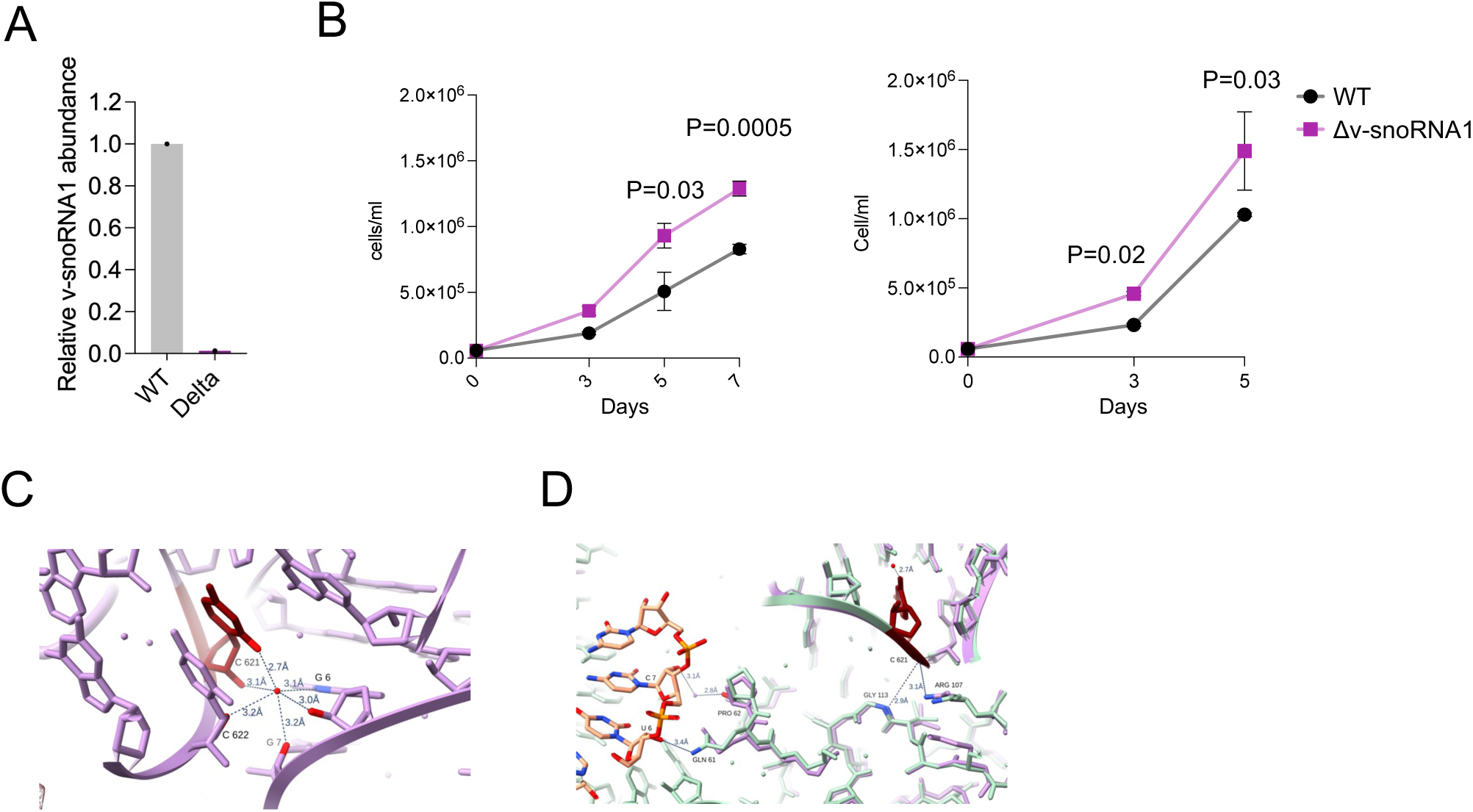
Infection of primary B-cells with wild-type (WT) or mutant (Δv-snoRNA1) EBV M81. Related to. Figure 5. (A) qPCR validation of loss of v-snoRNA1 expression in Δv-snoRNA1-infected LCLs. (B) Proliferation of LCLs from two additional donors showing increased proliferation of Δv-snoRNA1-infected cells relative to WT-infected cells. Two biological repeats were performed for each donor. (C) A network of contacts of the water molecule formed with neighboring 18S residues in ribosome structures of polysomes purified from LCLs infected with Δv-snoRNA1 of the M81 EBV strain. Purple – structure in Δv-snoRNA1-infected cells, red- C621. (D) C621 interacts directly with the backbone amide of Gly113 and the side chain of Arg107 of uS12ribosomal protein (RPS23). Purple – structure in Δv-snoRNA1-infected cells, red- C621. Green and orange – SPI9 with mRNA (orange).

**Supplementary Figure 3.**
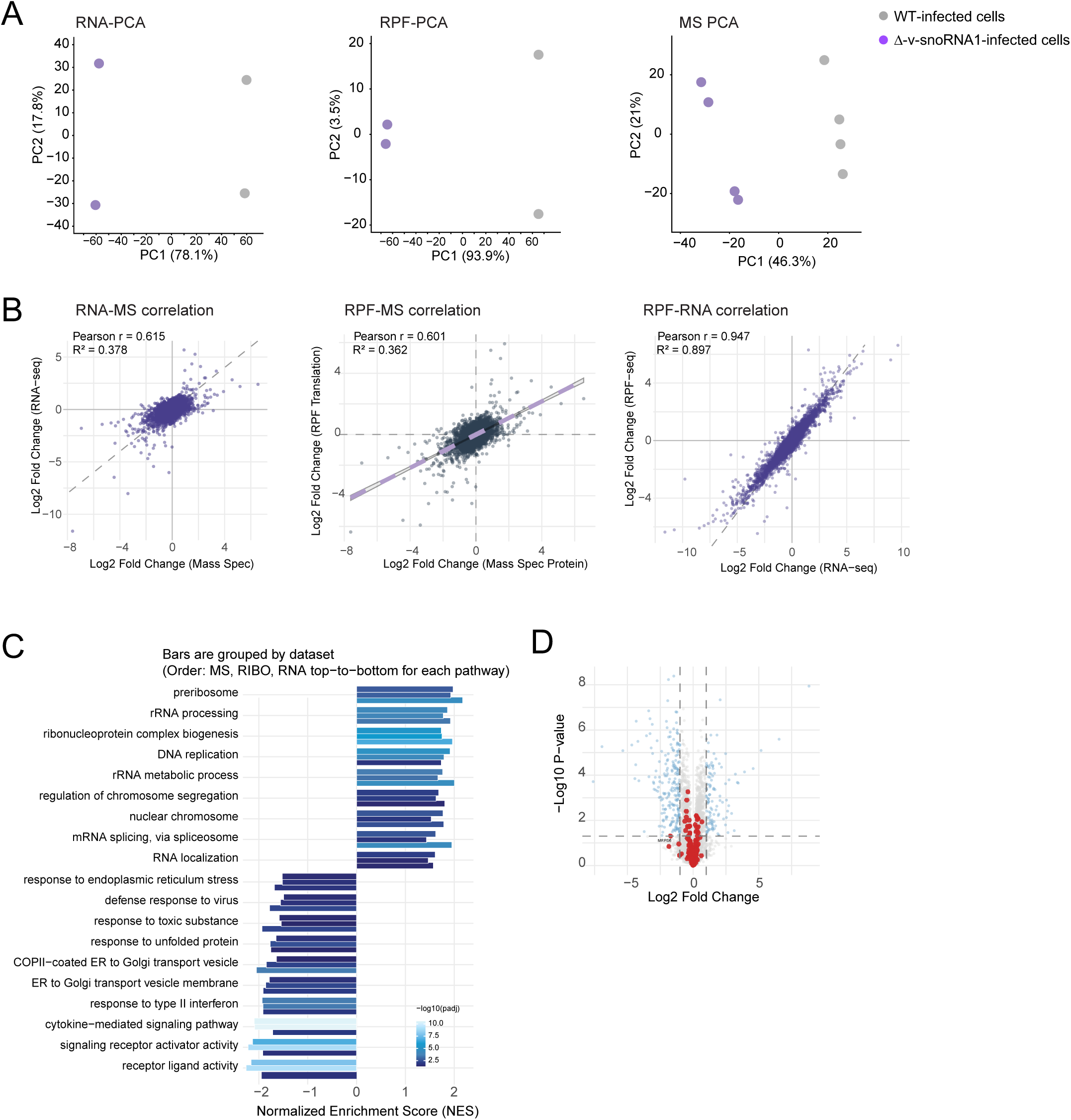
Ribosome and proteomic profiling of EBV WT- and Δv-snoRNA1-infected LCLs. Related to. Figure 6. (A) PCA of RNA-seq, RFP and proteomics of WT- and Δv-snoRNA1-infected samples, showing a clear discrimination of samples. WT-infected sample – grey, Δv-snoRNA1-infected samples – purple. (B) Correlations of the various data sets. (C) Commonly enriched pathways in RNA-seq, RPF and proteomics, showing each enrichment’s p-adjust. In each pathway, the order of the datasets is MS-RFP-RNA from top to bottom. (D) Volcano plot of differential abundance of ribosomal proteins in WT- and Δv-snoRNA1-infected cells. No significant differences were found.

## 1. EXPERIMENTAL MODEL AND STUDY PARTICIPANT DETAILS

### Cell Lines

Human HeLa cells were cultured in Dulbecco’s Modified Eagle’s Medium (DMEM) supplemented with 10% fetal bovine serum (FBS) and 100 µg/mL penicillin–streptomycin. All other cell lines were maintained in RPMI 1640 supplemented with 10% FBS, 100 µg/mL penicillin–streptomycin, 1% L-glutamine, 1% sodium pyruvate, and 1% non-essential amino acids. Lymphoblastoid cell lines (LCLs) carrying wild-type (WT) M81 or Δv-snoRNA1 M81 Epstein–Barr virus strains were kindly provided by Henri-Jacques Delecluse (DKFZ, Heidelberg, Germany).

## 2. METHOD DETAILS

### Plasmid Construction and Lentiviral Transduction

For construction of the pIRESpuro/GL/v-snoRNA1 expression plasmid, the coding region of the EBV *v-snoRNA1* gene was PCR-amplified using genomic DNA purified from an EBV-positive human B lymphoblastoid cell line TK6 (ATCC; CRL-8015) as template and inserted into the *Cla*I and *Xho*I sites of the pIRESpuro/GL expression plasmid^24^ by using the PCR-introduced *Cla*I and *Xho*I restriction sites as described earlier^24^. The *v-snoRNA1* gene was inserted into the second intron of a truncated globin gene under a CMV promoter. For lentiviral expression of v-snoRNA1, v-snoRNA1m and snoRD127, used for transduction of BJAB cells, annealed oligodeoxynucleotides spanning the full-length of the relevant snoRNA gene were cloned either into the pLKO.1 plasmid (Addgene #10878, a gift from David Root), or into pLKO.5 (used for puromycin incorporation assays) which contains a GFP cassette (Addgene #57822 was a gift from Benjamin Ebert). When possible, transduced cells were selected by puromycin, alternatively GFP+ cells were sorted by FACS. For viral production, 293T cells were transfected with the relevant lentiviral plasmid together with the packaging plasmids pM2.G (Addgene #12259) and psPAX2 (Addgene #12260) (which were gifts from Didier Trono).

### RNA isolation and analyses

Cellular RNAs from cultured cells were isolated by the guanidinium thiocyanate-phenol-chloroform extraction procedure^32^. For northern blot analyses of precursor and mature rRNAs, 4 µg of total RNA was denatured by incubation in NorthernMax™-Gly Sample Loading Dye (Invitrogen) for one hour at 55°C before being size-fractionated on a 1.2% agarose gel containing 10 mM PIPES, 30 mM BIS-TRIS and 1 mM EDTA. To facilitate RNA transfer onto a NylonN+membrane (GE Healthcare), the separated RNAs were partially fragmented in situ by incubation of the gel in 75 mM NaOH for 20 min. The gel was neutralized by washing in 0.5 M Tris-HCl, pH 7.4, 1.5 M NaCl solution two times for 15 min. The neutralized gel was incubated in 10 x SSC for 20 min, before being subjected to passive overnight RNA transfer in 10 x SSC. The RNAs were cross-linked to the membrane by irradiation with 254 nm UV light and detected with hybridization of 5’-terminally radiolabeled sequence-specific oligodeoxynucleotide probes specific for ITS1, ITS2, 18S, 28S and 5.8S rRNAs. Radioactive signals were revealed by using an intensifying screen and Typhoon Trio PhosphoImager (GE Healthcare).

Detection of RNAs by RNase A/T1 protection and mapping of 2’-O-methylated ribonucleotides by primer extension in low dNTP concentration have been described^32^. For identification of 2’-O-methylated nucleotides resistant to alkaline hydrolysis, total cellular RNA (10 µg) was partially hydrolyzed in 20 µl of 80 % formamide containing 0.4 mM of MgCl_2_ at 100 °C. After 5, 10, 15 and 20 min of incubation, 5 µl samples were collected and combined on ice. RNA was precipitated with ethanol and used as template for primer extension analysis with the Superscript II reverse transcriptase as recommended by the manufacturer (Invitrogen).

### Measure of nascent protein synthesis

About 1 million of exponentially growing cells were incubated in the presence of 50 µM Click-iT® homopropargylglycine (Invitrogen) for 1 hour. Cells were harvested by trypsin dissociation, washed in PBS and fixed with 4% paraformaldehyde in PBS for 15 min. The fixed cells were permeabilized with 0.25% Triton X-100 in PBS for 15 min and washed twice with in PBS containing 3% bovine serum albumin (BSA). To chemoselectively ligate Alexa Fluor 488 Azide (Invitrogen) to incorporated homopropargylglycine, we used Invitrogen’s Click-iT™ Cell Reaction Buffer Kit according to the manufacturer’s instructions. After ligation for 30 min, the excess of Alexa Fluor 488 Azide was removed by rinsing with PBS containing 3% BSA. The cellular fluorescence was measured by flow cytometry on a CytoFLEX S Flow Cytometer (Beckman Coulter).

### SunRiSE

50,000 HeLa or BJAB cells per well were plated in 24well plate or a 96U plate, respectively, a day prior to experiment. Cells were treated with 5ug/ml harringtonine and puromycin was added at the indicated time points at 3ug/ml. Cells were washed twice with cold PBS, harvested, and then were fixed and permeabilized using BD CytofixCytoperm kit (cat BD555028), according to manufacturers’ instructions. Staining wash was done in Perm/Wash buffer, supplemented with 2% human serum (for blocking) at a dilution of 1:500. Anti-puromycin antibody was the clone 12D10, purchased from Sigma-Aldrich. Secondary antibody was Rabbit anti-Mouse IgG (H+L) Cross-Adsorbed Secondary Antibody, Alexa Fluor 647 (Thermo Fisher Scientific, cat A21239).

### Sucrose gradient

The culture medium of 10 million exponentially growing cells was supplemented with 100 µg/ml cycloheximide, and cells were further grown for 20 min before being harvested in PBS supplemented with 100 µg/ml cycloheximide. Cells were collected by centrifugation at 300 x g for 5 min and resuspended in 1 ml of Hypotonic Buffer (10 mM HEPES pH 7.9, 10 mM KCl, 1.5 mM MgCl_2_, 100 µg/ml cycloheximide). After incubation on ice for 10 min, cells were disrupted in a Dounce homogenizer by 20 strokes with pestle B. Cytoplasmic extract was clarified by 5 min centrifugation at 600 x g at 4 °C. The supernatant was further purified by two steps of centrifugation at 12,000 x g, for 5 min at 4 °C. Protein content of the cytoplasmic extracts was measured by Bradford reagent and extract samples containing 0.8 mg protein were loaded onto 10% - 50% sucrose gradients prepared in Hypotonic Buffer. The gradients were centrifuged in a SW41 rotor at 39 K rpm, at 4 °C for 2 hours and stopped without break in a Beckman Coulter ultracentrifuge. Ribosome distributions along the gradients were measured by UV absorbance monitoring with a Teledyne ISCO fraction collector.

### Dual luciferase mistranslation reporter assay

Cells plated on 12-well plates were transfected with the pRM hRluc-hFlucD357X(TAG), pRM hRluc-hFlucD357X(TGA), pRM hRluc-hFlucH245R(CGC) or pRM hRluc-hFlucH245R(AGA) expression vectors using JetPRIME (OZYME Polyplus) transfection reagent^78^. After 36 hours of growth, the transfected cells were collected and lysed in 100 µl of Cellytic M buffer (Sigma-Aldrich). For detection of both Renilla and firefly luciferase activities in a luminometer, we used 30µl of clarified cell extracts and 20µl of Renilla or Firefly Luciferase Assay reagent (Promega).

### Cell cycle analysis

Cells were cultured a day before at equal density of 3.5×10⁵ cells/ml. Day of the experiment, cells were washed and fixed with cold 70% ethanol, then treated with RNAse A of 0.1 mg/ml final concentration, and with propidium iodide (PI) of 50 µg/ml final concentration. PI emission was measured by FACS.

### RiboMethSeq and computational analysis

0.2-2 ug total RNA was fragmented by alkaline hydrolysis in a final concentration of 50mM bicarbonate buffer and heated to 95°C for 10 minutes. To stop the fragmentation, 1ml of pre-cooled participation mix (1ml absolute ethanol, 1% 3M NaOAc, and 1% glycogen per reaction) was added, and samples were flash frozen in liquid nitrogen. The mix was centrifuged for 30 min at 4°C and subsequently washed with 80% ethanol. End repair was performed using Antarctic phosphatase and T4 PNK prior to library preparation by NEBNext small RNA library kit. At least 10M reads were obtained for each library. We performed quality trimming and adapter clipping of the obtained reads using Trimmomatic (v0.39). In RiboMethSeq analysis, only short reads with properly clipped adapters at the 3’-end were retained. Next, we mapped reads to human reference rRNA sequences (18S, 28S, 5S, and 5.8S) using Bowtie2, keeping only uniquely mapped reads. Both 5’- and 3’-ends of the fragment were utilized. Methylation levels were quantified by calculating the MethScore for each rRNA position.

### Ribosome Profiling and computational analysis

Cells were treated with cycloheximide (100 µg/ml, 15 min) to stabilize translating ribosomes and lysed on ice in polysome lysis buffer (20mM Tris pH7.4, 5mM MgCl_2_, 150mM NaCl, 1%Triton, supplemented with 1% deoxycholate, 100 µg/ml cycloheximide, 1 mM DTT, murine RNase inhibitor, protease inhibitors, and DNase I). RNA concentrations were determined by Nano-drop. 10% of each lysate was reserved for total RNA sequencing, which was performed in parallel using the KAPA mRNA HyperPrep Kit (Roche). Ribosome-protected fragments (RPFs) were generated by micrococcal nuclease digestion (20 min, 25°C) and purified by ultracentrifugation through a 34% sucrose cushion (TLA120.2 rotor, 111,000 rpm, 1h, 4°C). Ribosomal pellets were resuspended in lysis buffer, and RNA was extracted using a column-based kit. RPFs of 26-34 nucleotides were size-selected on 15% TBE–Urea polyacrylamide gels, visualized by SYBR Gold staining, excised, and eluted. Purified fragments were dephosphorylated with Antarctic phosphatase and rephosphorylated using T4 polynucleotide kinase. Libraries were prepared with the NEBNext Small RNA Library Prep Kit (NEB) according to the manufacturer’s instructions. For both ribosome protected fragments (RPF)-sequencing and RNA-seq, initial quality control was conducted using FastQC (0.11.9). Then, adapter trimming and length filtering were performed with Cutadapt (3.5). Reads were aligned to the human genome version hg38 and to the M81 genome reference using STAR (2.7.10a). For the RPF-seq data, only aligned reads with length of 24-36 nucleotides were retained using SAMtools (1.16.1). Gene expression counts were obtained using HTSeq (0.13.5) with the ‘union’ mode, specifying ‘CDS’ for ribo-seq and ‘exon’ for RNA-seq. RNA-seq data were normalized to fragments per kilobase of transcript per million mapped reads (FPKM) using transcript length, while RPF-seq data were normalized to coding sequence (CDS) length. Genes with log transcript per million (TPM) counts below 4 in all conditions were filtered out, and differential expression analysis was performed using DESeq2 with false discovery rate (FDR) correction. Translation efficiency (TE) was calculated by dividing ribosome-protected fragment (RPF) abundance by mRNA abundance for each gene. Log₂-transformed FPKM values from RPF-seq and RNA-seq derive a TE score that reflects ribosome occupancy per transcript.

### Ribosome purification for structural analysis

Cells were resuspended in ribosome lysis buffer (30 mM Tris pH 7.5, 20 mM Mg acetate, 300mM NaCl, 2% tritonX-100, 40U/ml cells RNAsin (Promega), Complete mini protease inhibitor) and lysed by inverting gently. Lysate centrifuged at 9500 rpm for 10 min at 4 °C. Supernatant was loaded onto a sucrose cushion (45mM HEPES KOH pH=7.5, 150mM K acetate, 10mM Mg acetate, 2mM DTT, 1.1M sucrose. Buffer was treated with a 5mg of bentonite clay and filtered before use) and centrifuged overnight at 45000 RPM, 4°C. Pellet was resuspended in a resuspension buffer (20mM Tris pH 7.5, 150mM K acetate, 10mM Mg acetate, 1mM DTT, 40U/ml RNAsin) and loaded onto sucrose gradients (10–40% sucrose, 20mM, Tris pH 7.5, 150mM K acetate, 10mM Mg acetate, 1mM DTT. Buffer was treated with a 5mg of bentonite clay and filtered before use) and centrifuged at 22,000rpm for 11h at 4°C. Sucrose fractions were collected and pelleted by centrifugation at 45,000rpm overnight at 4°C. Pellets were resuspended in a final buffer (20mM Tris pH 7.5, 100mM K acetate, 10mM Mg acetate, 10mM NH_4_Cl, 1mM DTT) and stored at −80°C. The resulting 80S and polysome concentrations were 25A and 70A for WT, and 100A and 150A for the mutant, respectively.

### Cryo-EM data collection and refinement

3 µl of purified polysomes of each sample were applied onto freshly glow-discharged holey carbon grids (Quantifoil R2/2) coated with a continuous carbon film, and incubated for 30 s at 4°C, 100 % humidity in a Vitrobot Mark IV (Thermo Fischer Scientific). The grids were blotted for 2.5 s before plunge freezing in liquid ethane. Micrographs were acquired using a Titan Krios electron microscope (Thermo Fischer Scientific) operated at 300 kV. Images were acquired using K3 direct electron detector (Gatan Inc.). Magnification was 105,000, with a pixel size of 0.8242 Å/pixel and a dose rate of ∼1 electron/Å^2^/s. Defocus values ranged from −0.5 to −1.5 μm. Relion 5.0.0 was used for data processing. Motion correction was performed using Motioncor2. The contrast transfer function parameters were estimated using CTFFIND-4.1. Referenced particle picking resulted in a particle count of 6,553,806 and 859,381 for mutant and WT, respectively. 1,709,343 and 316,464 particles were used for 3D classification (mutant and WT, respectively). Classes appearing to have well-formed 80S particles were selected. A total of 386,469 and 178,121 particles (mutant and WT, respectively) were used for auto-refinement. Following refinement, particles were subjected to CTF refinement, particle polishing, and 3D refinement to calculate 2.1 Å and 2.2 Å EM maps for mutant and WT, respectively. The resulting 3D EM maps were then subjected to a cycle of multibody refinement using separate masks for different ribosome parts. The large subunit (LSU), the small subunit (SSU) body and head, produced density maps at 2.02 Å, 2.15 Å, and 2.28 Å resolution for Mut, and 2.1 Å, 2.28 Å, and 2.48 Å resolution for WT, respectively. The gold standard Fourier shell correlation (FSC) value criterion of 0.143 was used to determine average map resolutions, as implemented in Relion 5.0.0.

### Model Building and Refinement

Model building was executed using a template-guided method, using the previously published structure 8QOI. The starting model was docked into the maps using UCSF ChimeraX 1.1 and manually adjusted using Coot 0.9.6. Model refinement was performed using an iterative approach, including real-space refinement and geometry regularization in Coot. Structural analysis and figures were generated using Coot 0.9.6 and UCSF ChimeraX 1.1.

### Network Centrality and Master Regulator Analysis

To identify putative master regulators of the transcriptional response, we performed a network centrality analysis by integrating the differential expression results with the DoRothEA (Discriminant Regulon Expression Analysis) database. A transcription factor (TF)-target interaction network was constructed utilizing high-confidence human regulons (Confidence levels A and B). The network was subset to include only interactions where target genes reached statistical significance (FDR < 0.05). Using the igraph R package, a directed graph was modeled, and hub centrality scores were calculated via Kleinberg’s hub centrality algorithm to quantify the influence of each TF over the observed differential expression signature. TFs were ranked by their hub scores, with those approaching a centrality value of 1.0 identified as key regulatory hubs coordinating the transition from the wild-type to the knockout state. Significant TFs were further characterized by their Log2 Fold Change to distinguish between activators and repressors associated with the knockout phenotype.

## 3. QUANTIFICATION AND STATISTICAL ANALYSIS

All statistical analyses were performed using R (v. 3.6.0). Statistical tests used for each experiment with details on sample size (n) are indicated in the corresponding figure legends. For comparisons between two groups, unpaired two-tailed Student’s t-tests were used unless otherwise specified. A p value < 0.05 was considered statistically significant. For high-throughput datasets (RNA-seq, Ribo-seq, and RiboMethSeq), adjusted p values were calculated using the Benjamini–Hochberg method to control the false discovery rate (FDR), and adjusted p value < 0.05 was used as the significance threshold. Statistical analysis of differential translation analysis is detailed in figure legends.

## 4. KEY RESOURCES TABLE

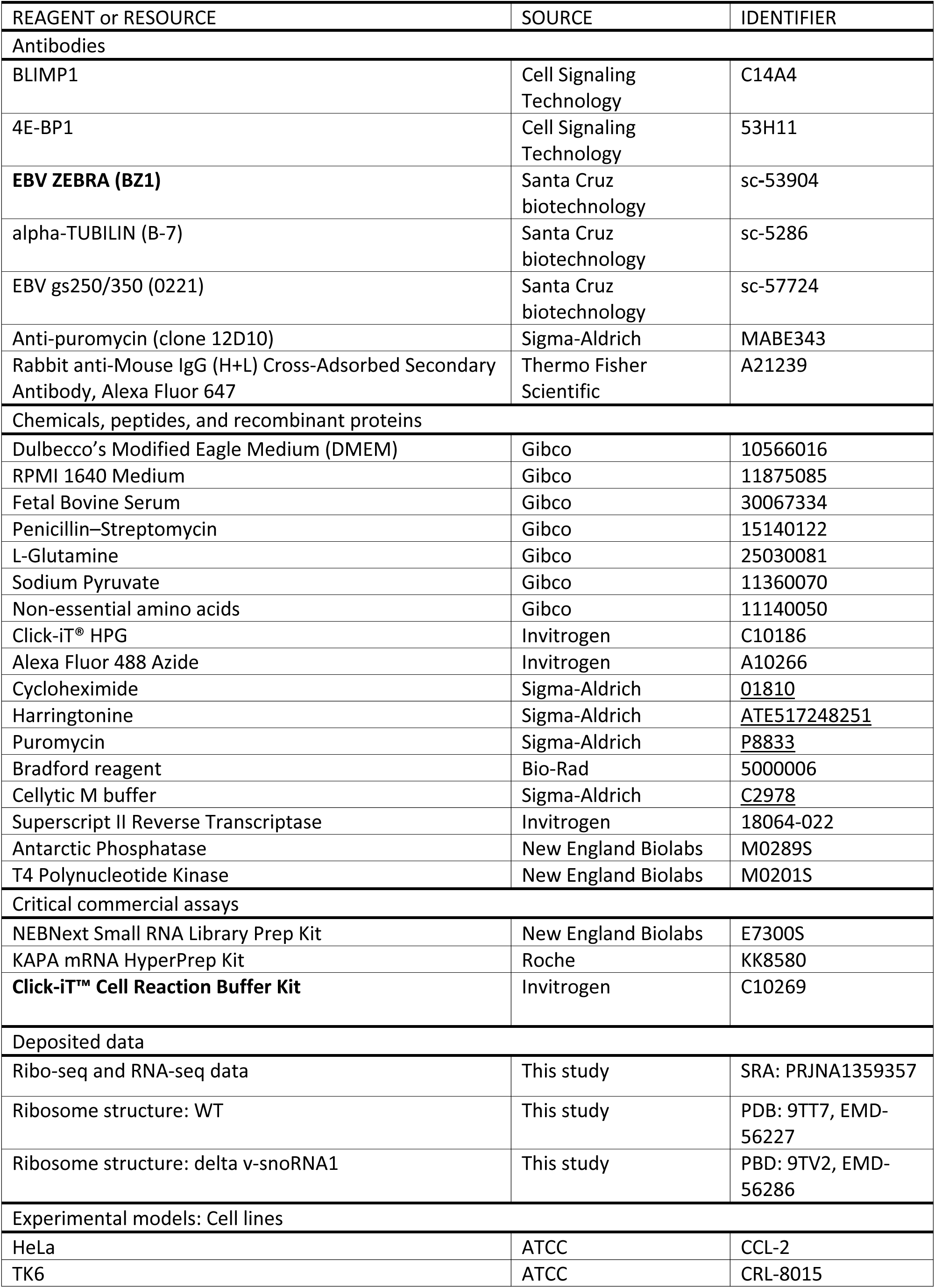
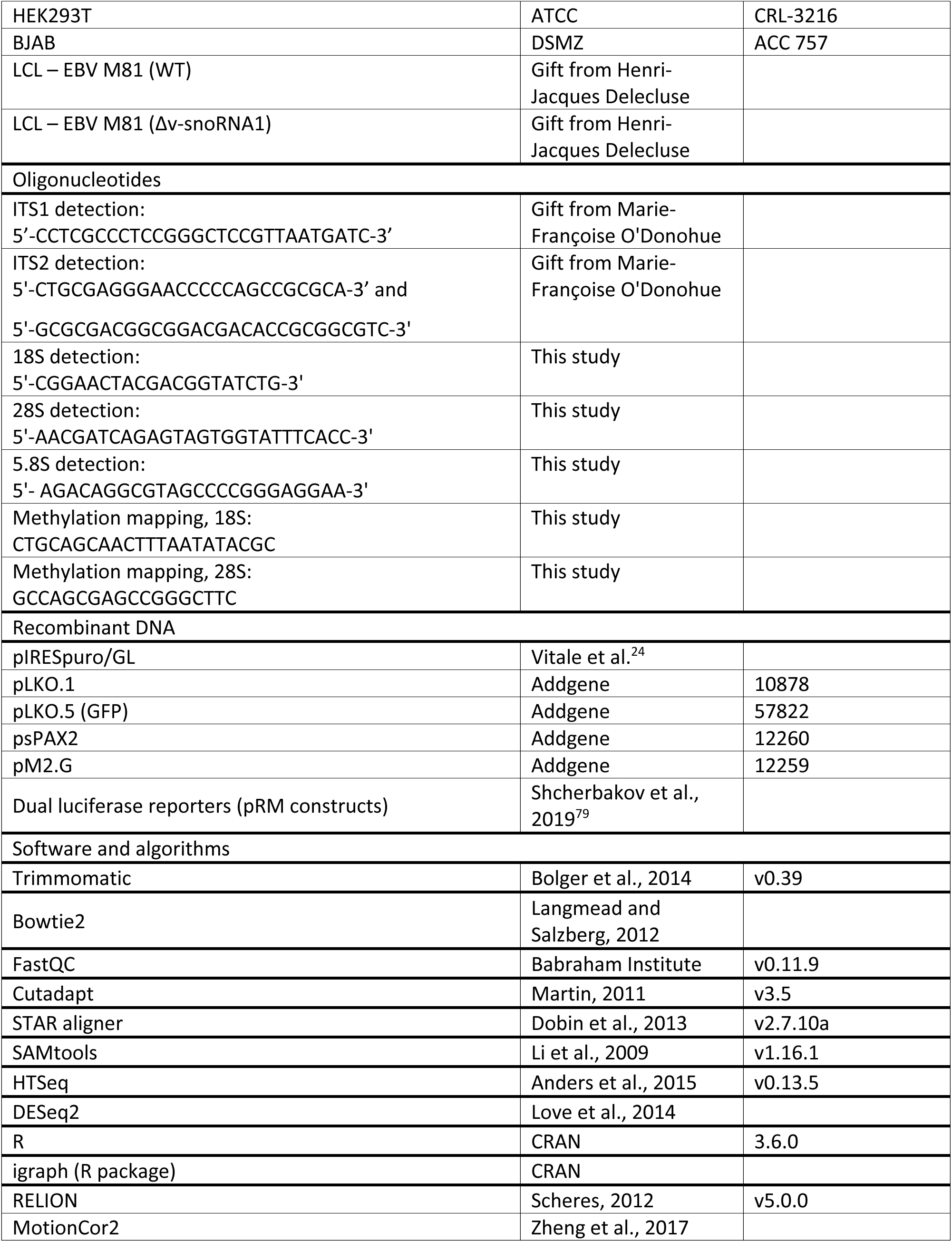
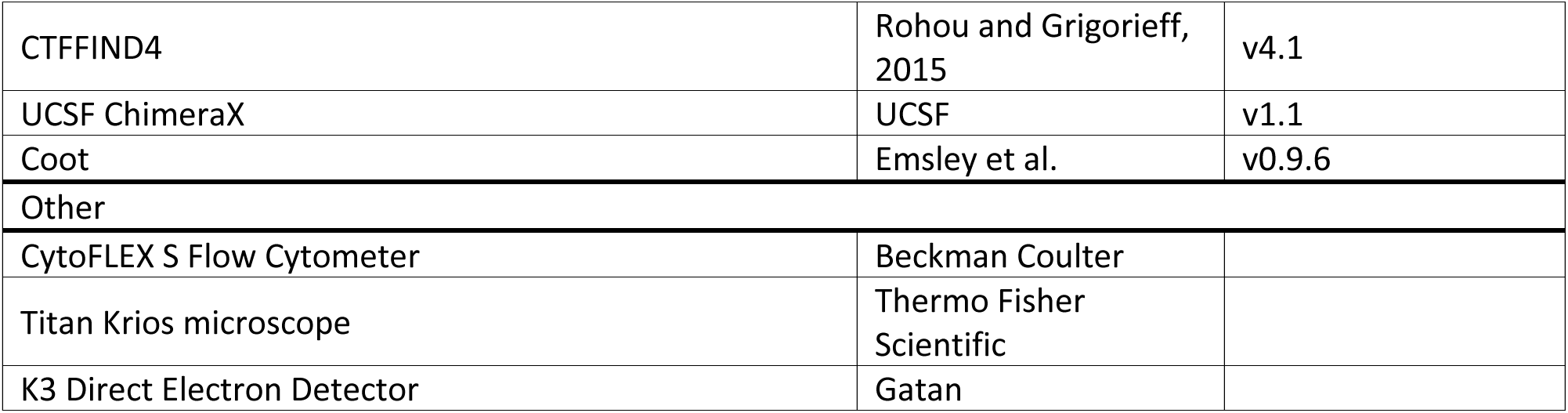

## Notes

### Competing Interest Statement

The authors have declared no competing interest.

